# Extraocular Photoreception Controls Nocturnal Hatching via a Neuroendocrine Pathway in False Clownfish

**DOI:** 10.64898/2026.09.14.750986

**Authors:** Sakuto Yamanaka, Takahiro Yamashita, Yasuhiro Kamei, Hiroyuki Takeda, Takashi Yamanaka, Masato Kinoshita

**Affiliations:** Faculty of Life Sciences, Kyoto Sangyo University Motoyama, Kamigamo, Kita-ku, Kyoto 603-8555, Japan; Department of Biophysics, Graduate School of Science, Kyoto University Kitashirakawa Oiwake-cho, Sakyo-ku, Kyoto 606-8502, Japan; Trans-Scale Biology Center, National Institute for Basic Biology 38 Nishigonaka, Myodaiji, Okazaki, Aichi 444-8585, Japan; Division of Applied Biosciences, Graduate School of Agriculture, Kyoto University Kitashirakawa Oiwake-cho, Sakyo-ku, Kyoto 606-8502, Japan

## Abstract

Hatching is the transition from embryo to larva in oviparous animals. Birds, reptiles, and insects rupture the eggshell mechanically, whereas fish and amphibians digest the egg envelope using hatching enzymes. When and where larvae emerge influences their fitness. Many species regulate hatching timing using environmental cues, such as light, oxygen availability, and physical stimuli. In fish, hatching is driven by a neuroendocrine event that triggers hatching enzyme secretion; however, the mechanisms controlling this event are unknown. Using the false clownfish *Amphiprion ocellaris*, which hatches synchronously within 1–2 h after sunset, we showed that the role of thyrotropin-releasing hormone (TRH) as a hatching hormone, previously established in zebrafish and medaka, extended to clownfish, where TRH activated Ca²⁺ signaling in hatching gland cells through its receptor *trhra*, reinforcing the TRH–Ca²⁺–hatching enzyme axis as a conserved core module of teleost hatching. In clownfish, this module was additionally placed under upstream control by extraocular photoreception because dark-induced hatching persisted in eyeless embryos. This photoreceptive system was most sensitive to green light (550 nm), and several opsins with absorbance maxima in the corresponding middle-to long-wavelength range were expressed outside the eyes of pre-hatching embryos. Coupling extraocular photoreception to a conserved neuroendocrine module placed a critical life-history transition under stringent photic control, releasing larvae only in complete darkness, an adaptation that may have underpinned the success of this lineage in predator-rich coral reefs.

## Introduction

Teleosts have evolved hatching strategies adapted to their environment that are critical for the survival of embryos and/or larvae. The timing and environmental context of larval hatching can significantly affect subsequent survival, whether hatching occurs when food is abundant, when predation risk is low, or when larvae can be transported to appropriate nurseries. In many fish species, hatching is regulated by environmental cues including light, hypoxia, and water flow, making it one of the earliest environmentally responsive events in their life history. For example, California grunion (*Leuresthes tenuis*) and mummichog (*Fundulus heteroclitus*) lay their eggs in coastal sand, where embryos develop and wait to hatch until exposed to seawater during high tides (Dimichele & Taylor, 1980, 1981; Griem & Martin, 2000; Martin et al., 2011; Smyder & Martin, 2002). Light also has a major influence on the timing of hatching; medaka tend to hatch during the daytime (Yamagami, 1988), whereas species such as halibut (*Hippoglossus hippoglossus*) and false clownfish (*Amphiprion ocellaris*) hatch at night (Forsell et al., 1997; Helvik & Walther, 1992, 1993; McAlary & McFarland, 1993). The molecular mechanisms regulating hatching involve hatching enzymes that digest the inner layer of the chorion immediately before hatching (Kawaguchi et al., 2013; Yasumasu et al., 1989) and calcium signaling that regulates their secretion from hatching gland cells (Schoots et al., 1981). In addition, thyrotropin-releasing hormone (TRH) directly activates hatching gland cells, thereby functioning as a hatching hormone in zebrafish and medaka (Gajbhiye et al., 2024), although it is classically known as a hypothalamic hormone that acts on the pituitary gland as part of the hypothalamic–pituitary–thyroid axis. Although the internal molecular mechanisms regulating hatching have become increasingly understood, how these mechanisms integrate with environmental information remains largely unknown because of the weak link between hatching and environmental cues in model organisms such as zebrafish. The false clownfish (*Amphiprion ocellaris*) inhabits coral reefs and spawns adhesive eggs on rocks or coral surfaces. Embryos hatch synchronously within 1–2 h after sunset on a specific hatching day, namely 8 days post-fertilization (dpf) (Salis et al., 2021; Fobert et al., 2019, 2021; Yamanaka et al., 2021, 2025). Hatching occurs under strict and robust light regulation, such that even minimal light exposure can strongly suppress hatching (Fobert et al., 2019; Yamanaka et al., 2021, 2025). In this study, using calcium imaging, gene knockout approaches, and spectral analysis, we revealed that the TRH– Ca²⁺–hatching enzyme system was conserved in false clownfish and that this system was regulated upstream by extraocular photoreception. Furthermore, spectral sensitivity analysis revealed that hatching showed high sensitivity to green light and that several opsins with absorbance maxima in the middle-wavelength (green) region are expressed in pre-hatching embryos.

## Results

### Light environments regulate conserved hatching system

Several species of Pomacentridae, including anemonefish, hatch at night (Gladstone, 2007; Kingsford, 1985; McAlary & McFarland, 1993); however, how embryos recognize environmental cues and trigger the hatching system, including the secretion of hatching enzymes, remains unclear. Our previous reports showed that light cues strongly suppressed hatching and continuous dark conditions induced chorion digestion in false clownfish (Yamanaka et al., 2021, 2025). In addition, TRH activates hatching gland cells via its receptor *trhra* to trigger the secretion of hatching enzymes in zebrafish and medaka (Gajbhiye et al., 2024). Therefore, we examined whether these molecules function as conserved downstream components of the nocturnal hatching pathway in false clownfish. First, *in situ* hybridization chain reaction (HCR) in clownfish embryos revealed that *trh* is expressed in the anterior ventral regions of the hypothalamus and that *trhra* is expressed in hatching gland cells located on the lateral surface of the posterior trunk (Fig. 1A, B). Expression of the other *trh* receptors, *trhrb* and *trhr2*, was not detected by RT-PCR (Fig. S1). We performed a chorion digestion assay to quantify the extent of chorion digestion as an indicator of hatching enzyme secretion. While chorion matrix proteins such as zona pellucida proteins (choriogenins) become highly cross-linked and insoluble after fertilization (Yasumasu et al., 2010), fragments digested by hatching enzymes become soluble and can be quantitatively measured using the bicinchoninic acid (BCA) assay, a protein quantification method, in sodium dodecyl sulfate (SDS) buffer. We collected eggs at 8 dpf, incubated them under light or dark conditions for 1 h, and then peeled off the egg envelope with fine tweezers to quantify the amount of digested chorion protein.

**Fig. 1.**
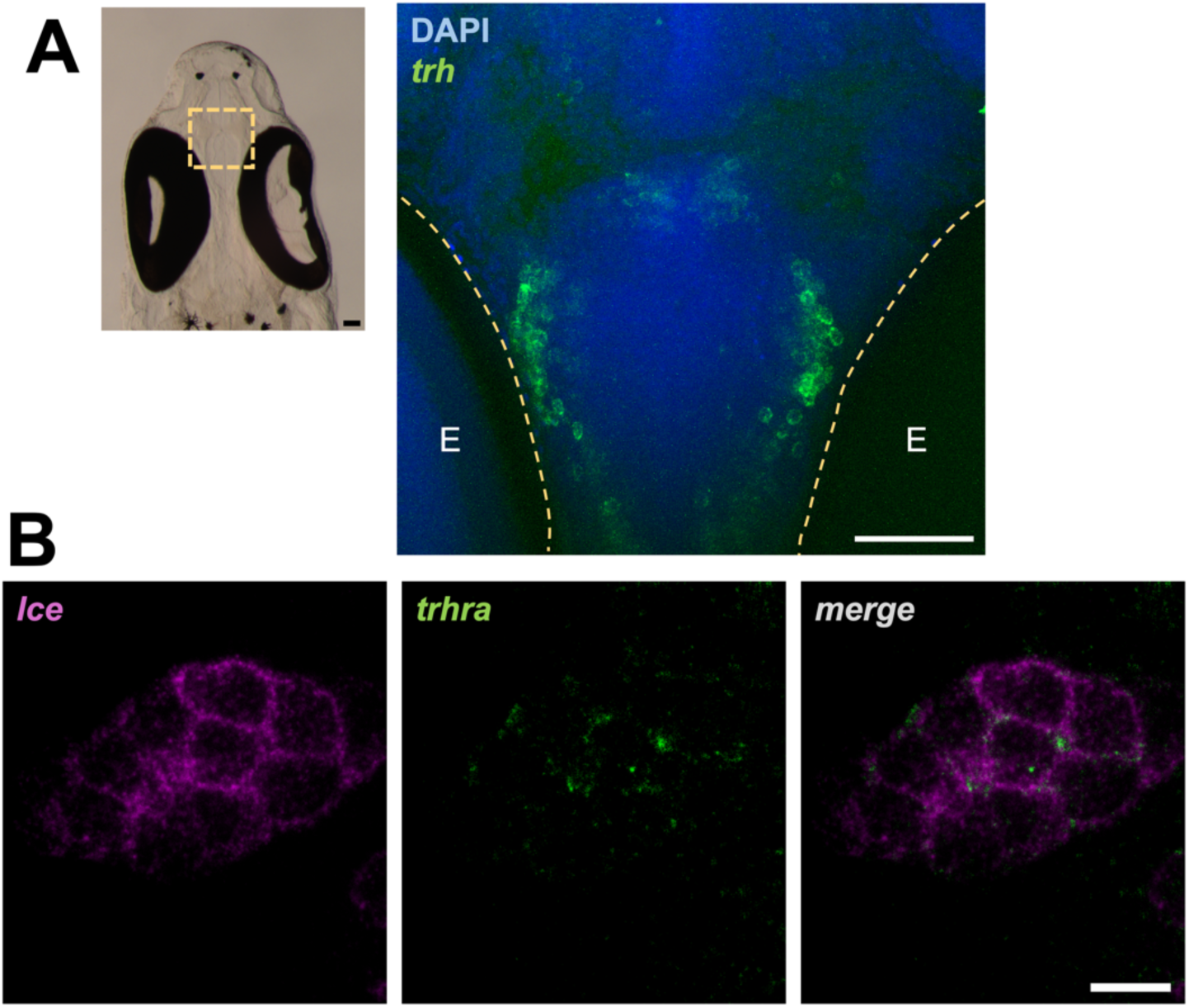
Expression patterns of *trh*. (A) and its receptor *trhra* (B). (A) The expression pattern of *trh* in the ventral side of hypothalamus. The bright-field image shows a horizontal vibratome section, and the fluorescent field shows a magnified view with HCR signal (blue, DAPI; green, *trh*). The white scale bar shows 50 μm. (B) The expression patterns of the *trhra* in hatching gland cells in the tail epidermis of a pre-hatching embryo (8 dpf). HCR signals (magenta, *lce*; green, *trhra*) show that *trhra* is specifically expressed in hatching gland cells. White scale bar shows 10 μm.

In wild-type embryos, dark conditions significantly induced chorion digestion, whereas light conditions did not (*p* = 1.6 × 10⁻⁴, Fig. 2A). Since false clownfish shows high CRISPR/Cas9 efficiency even in F0 embryos (Mitchell et al., 2021), we generated *trhra* mosaic knockout embryos using CRISPR/Cas9 and confirmed the high mutagenesis efficiency in these individuals using a heteroduplex mobility assay (Fig. S2). Under dark conditions, these embryos showed significantly less chorion digestion than the wild-type embryos (*p* = 0.0093, Fig. 2B). Furthermore, direct injection of TRH into the heart or tail of the embryos induced chorion digestion, even under light conditions, under which hatching did not occur normally (Fig. 2C). Chorion digestion differed significantly among groups (Kruskal–Wallis, *p* = 1.5 × 10⁻⁵). TRH-injected embryos under light (heart and tail) showed levels comparable to those of dark controls (Dunn’s test with Benjamini–Hochberg correction, *p* = 0.23 and 0.21, respectively), whereas Ringer-injected groups under light (heart and tail) remained significantly lower than those in the dark controls (*p* = 0.0001 and 0.0071, respectively). These results indicate that, similar to zebrafish and medaka, TRH acts as a hatching hormone that triggers the secretion of hatching enzymes in false clownfish. Furthermore, *Trhra* functions as a TRH receptor, suggesting that TRH secretion and its reception by *Trhra* are positioned downstream of photoregulation.

**Fig. 2.**
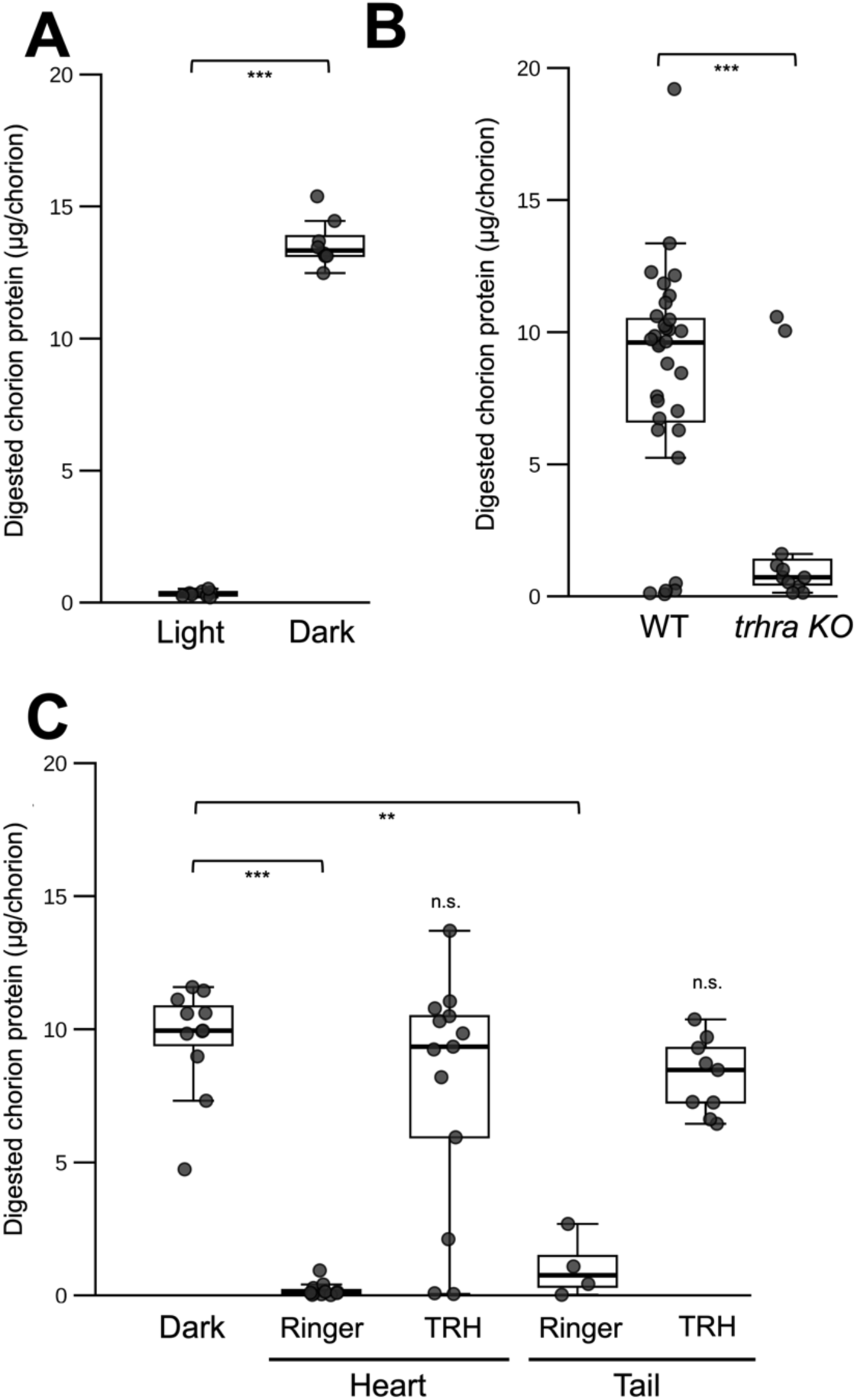
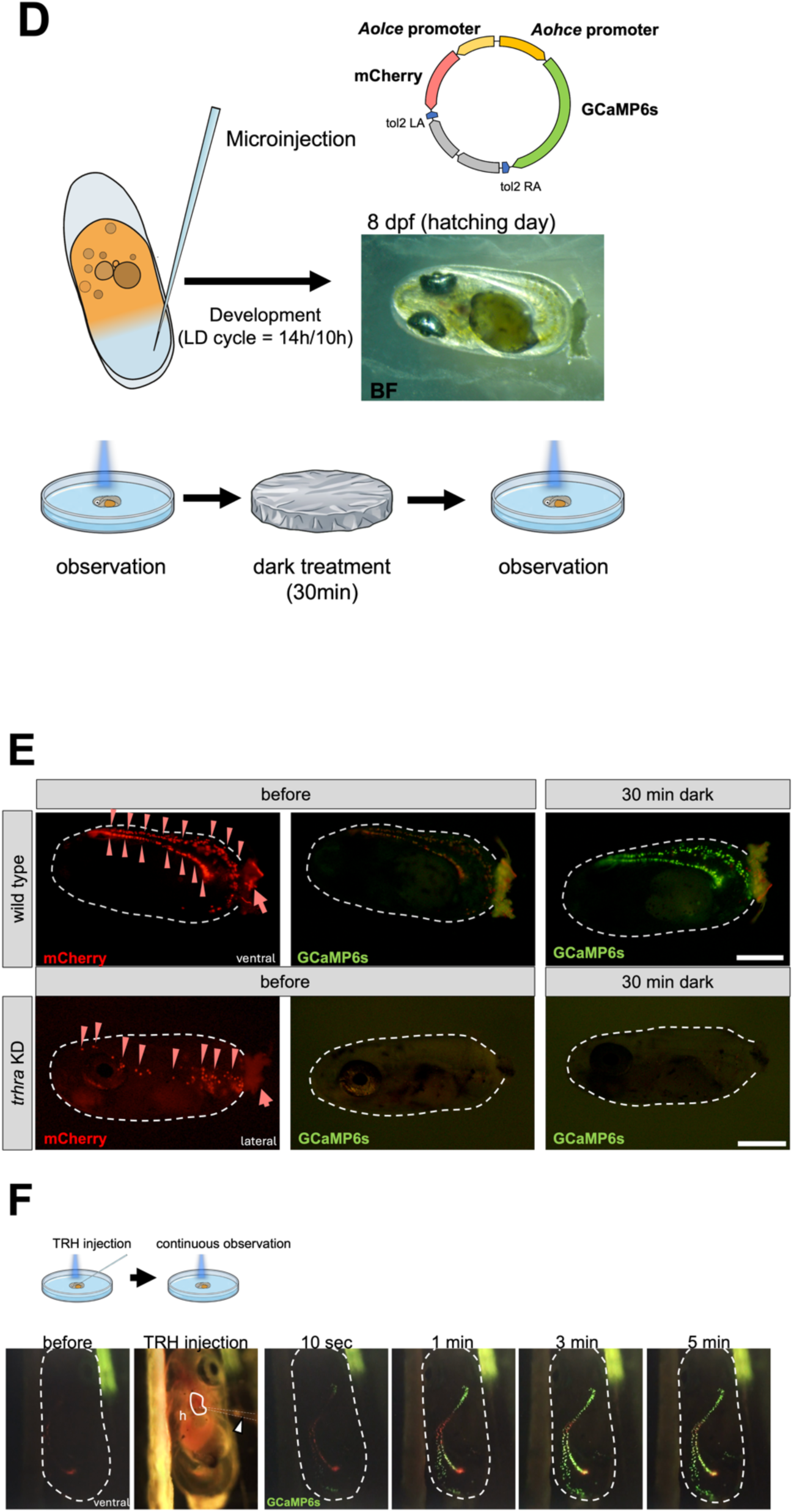
The extent of chorion digestion and Ca²⁺ imaging of hatching glands in *trhra* KO and TRH-injected embryos. (A) The extent of chorion digestion when pre-hatching embryos were incubated under light or dark conditions. Dark conditions induced significantly higher chorion digestion, but light conditions did not (Mann–Whitney U test, *p* = 1.6 × 10⁻⁴). *n* = 8 (light) and 8 (dark). (B) The extent of chorion digestion when wild-type or *trhra* knockout F0 embryos were incubated in the dark. Wild-type embryos showed high chorion digestion, but *trhra* mosaic KO embryos showed significantly suppressed chorion digestion (Mann– Whitney test, *p* = 0.0093). *n* = 16 (wild type) and 11 (*trhra* F0 crispant). (C) The extent of chorion digestion when 50 μM TRH solution or Ringer solution injected into the heart or tail muscle of embryos under light condition. Asterisks indicate significant differences from the dark control (Kruskal–Wallis followed by Dunn’s test with Benjamini–Hochberg correction; \*\**p* < 0.01, \*\*\**p* < 0.001; n.s., not significant). *n* = 11, 11, 9, 13 and 9 for the dark control, Ringer (heart), Ringer (tail), TRH (heart) and TRH (tail) groups, respectively. (D) A flowchart of calcium imaging using transgenic clownfish embryos. We generated mosaic transgenic clownfish for calcium imaging by microinjection with a reporter vector expressing GCaMP6s and mCherry driven by two hatching enzyme promoters (*hce* and *lce*), and incubated mCherry-positive embryos under an LD cycle (LD 14:10) for 8 days. On the day of hatching (8 dpf), green fluorescence was observed before and 30 min after complete dark exposure. (E) Calcium imaging in transgenic and *trhra* (a TRH receptor) mosaic knockout embryos. In embryos with intact *trhra*, the green fluorescence of GCaMP6s was specifically observed in mCherry-labeled hatching gland cells after 30 min of dark exposure (*n* = 3/3). In contrast, no GCaMP6s fluorescence was detected in *trhra* mosaic-knockout embryos under the same conditions (*n* = 0/2). These results show that dark exposure induces hatching gland activation via TRH and its receptor *trhra*. The white bars in each image show 0.5 mm. Arrowheads indicate labeled hatching gland cells. The arrows and dashed lines represent the adhesive filaments of the egg and egg envelope, respectively. (F) Calcium imaging of transgenic embryos after the cardiac injection of TRH. We microinjected the TRH solution into the embryo heart (h, white line) and continuously observed GCaMP6s fluorescence. Even under exposure to excitation light, strong fluorescence was detected in hatching gland cells within a minute after TRH injection (*n* = 2/2), indicating that TRH activated hatching gland cells independent of dark conditions. Dashed lines indicate the egg envelopes.

*Trhra* is a GPCR (G-protein-coupled receptor) that couples with *Gq* to trigger intracellular calcium signaling (Hinkle et al., 2012). Therefore, we performed calcium imaging of hatching gland cells using GCaMP6s (Chen et al., 2013) to visualize TRH-induced calcium activation. We generated mosaic transgenic clownfish for calcium imaging by microinjection of a reporter vector expressing GCaMP6s and mCherry driven by two hatching enzyme promoters (*hce* and *lce*) (Fig. 2D). After differentiation of the hatching glands, embryos exhibiting mCherry fluorescence in the hatching glands at 3 dpf were selected and reared under a normal light–dark (LD) cycle (LD 14:10) until the hatching day (8 dpf). The green fluorescence of GCaMP6s was not activated until the day of hatching, but strong fluorescence was observed after 30 min of continuous dark treatment in the afternoon of day 8 (*n* = 3/3, Fig. 2E). However, this calcium activation in the hatching glands by dark treatment was not observed in *trhra* knockout/GCaMP6s-transgenic embryos (*n* = 0/2, Fig. 2E). In addition, when TRH was injected into the hearts of embryos, the green fluorescence of GCaMP6s was detected within a minute even under excitation light (*n* = 2/2, Fig. 2F). These results indicate that continuous dark conditions trigger the secretion of TRH and that TRH immediately activates intracellular calcium signaling in hatching gland cells to induce the secretion of hatching enzymes. These results show that photoenvironmental cues affect the evolutionarily conserved hatching system (TRH-calcium signal – hatching enzyme) in false clownfish.

### Hatching occurs independently of ocular photoreception

In teleosts, the eyes are known to be the primary photoreceptive organs, and 8 dpf embryos of false clownfish show retinal pigment deposition and accumulation of guanine pigments in the reflective layer (Fig. 3A), indicating that the eyes are functional in detecting the light environment. To investigate the role of the eyes in the regulation of hatching, we disrupted *pax6b*, a teleost paralogue of the master eye-development regulator *pax6* (Kleinjan et al., 2008), and *strip1*, which is required for inner retinal development in zebrafish (Ahmed et al., 2022), using CRISPR/Cas9. In some mosaic-knockout embryos, the eyes were almost completely absent (Fig. 3A). Embryos that severely lacked eyes did not exhibit positive phototaxis after hatching and instead displayed circling behavior, and no hatching occurred before the hatching day. By using eye-lacking embryos with high mutation rates (Fig. S3), we examined whether hatching-enzyme secretion was induced by darkness without ocular light input. The effects of genotype and light conditions were assessed using a two-way Scheirer–Ray– Hare test. Light conditions had a strong effect on chorion digestion (*p* < 0.0001), whereas neither the genotype (*p* = 0.38) nor the genotype × light interaction (*p* = 0.70) were significant. The results showed that, similar to wild-type embryos, *pax6b/strip1 knock-out* embryos also exhibited chorion digestion induced by darkness and suppressed by light (Fig. 3B). These observations suggest that while the eyes of clownfish embryos may be functional at the time of hatching, the photoregulation of hatching does not rely exclusively on the eyes, indicating that extraocular photoreception contributes to the regulation of hatching enzyme secretion.

**Fig. 3.**
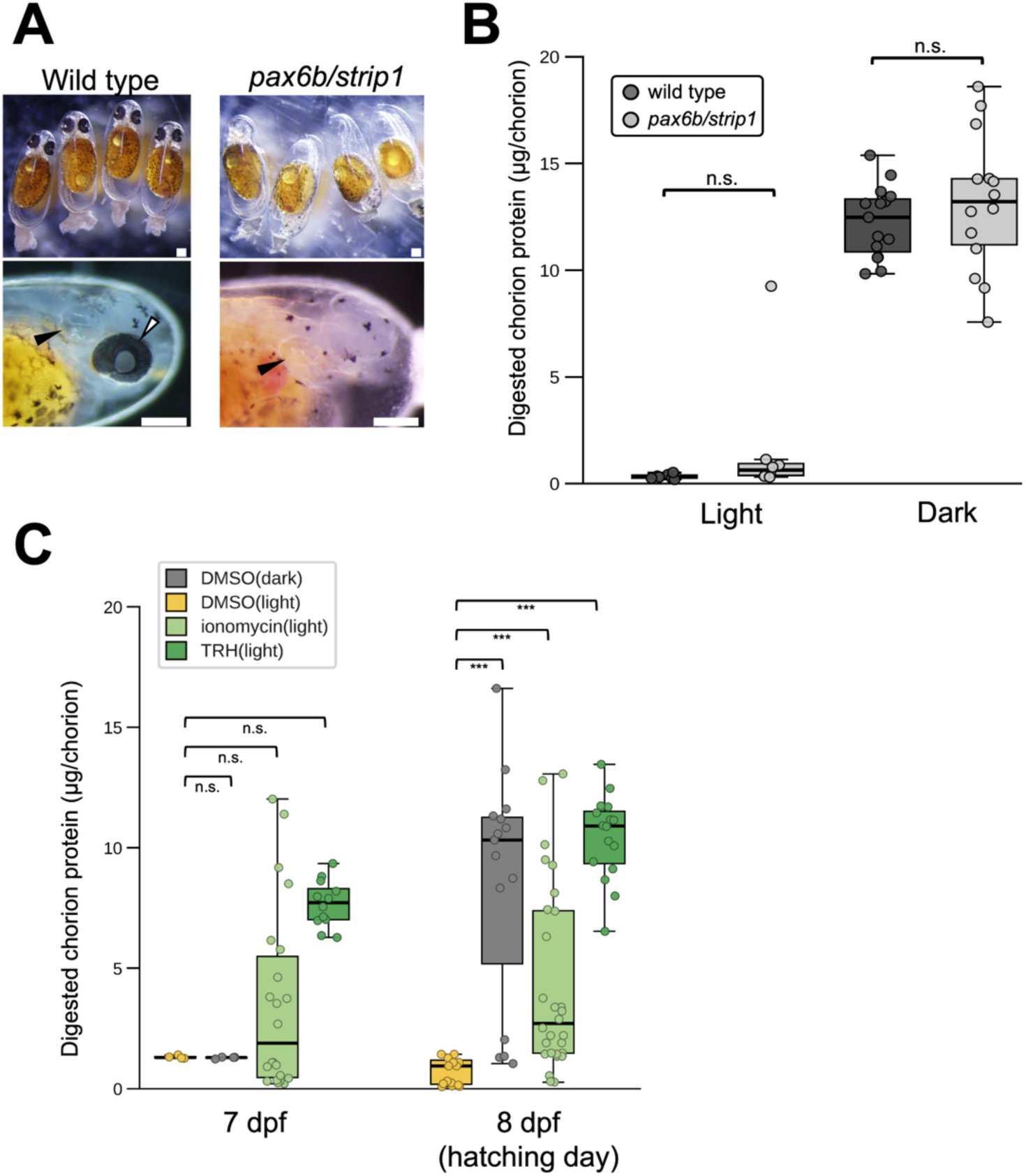
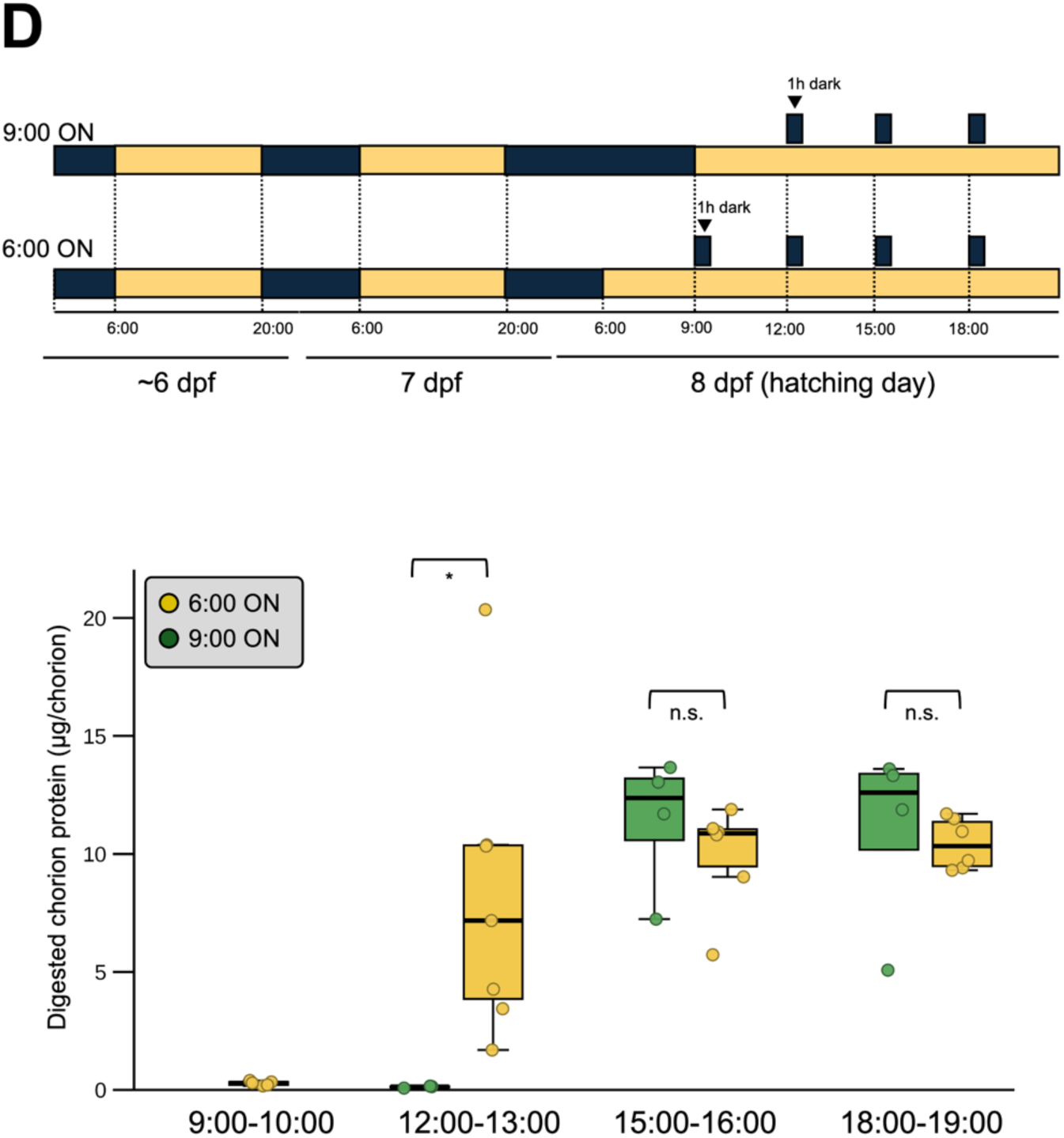
Chorion digestion in eyeless embryos and maturation of hatching glands. (A) Generation of eyeless embryos by CRISPR-mediated mosaic knockout of *pax6b* and *strip1*, regulators of eye and retinal development. Injected embryos showed varying degrees of eye loss, while body size was comparable to wild type. Black arrowheads show otic vesicle and white arrowhead shows eye. The white bars show 200 μ m. (B) Chorion digestion in eyeless embryos after 1-h light or dark exposure on the day of hatching (8 dpf). As in wild-type embryos, digestion was suppressed by light and promoted by darkness. Genotype had no significant effect (two-way Scheirer–Ray–Hare test, *p* = 0.38; interaction *p* = 0.70), whereas light condition did (*p* < 0.0001). Brackets, within-condition comparisons between genotypes (Mann–Whitney with BH correction). n.s., not significant. *n* = 8 (wild type, light), 15 (wild type, dark), 8 (eyeless, light) and 14 (eyeless, dark). (C) Effects of TRH and ionomycin on chorion digestion. Embryos were incubated for 1 h under light or dark conditions, or with TRH or ionomycin, at 7 and 8 dpf. Each treatment was compared with the light control of the same stage (Kruskal–Wallis followed by Dunn’s test with Benjamini–Hochberg correction; 7 dpf: *p* = 0.0023, *n* = 4, 4, 14 and 22 for light, dark, TRH and ionomycin; 8 dpf: p = 4.1 × 10⁻⁹, *n* = 15, 15, 16 and 28). Relative to the light control, all treatments increased chorion digestion at 8 dpf: darkness (*p* = 1.6 × 10⁻⁶), TRH (*p* = 4.3 × 10⁻⁹), and ionomycin (*p* = 5.8 × 10⁻⁴). At 7 dpf, darkness had no effect (*p* = 0.91). TRH raised the median to 7.72, narrowly missing significance after correction (*p* = 0.052), while ionomycin left the median largely unchanged (1.89, *p* = 0.91) but broadened the distribution. These results suggest that hatching gland cells may already be competent to secrete hatching enzyme in response to TRH before the day of hatching. (D) Chorion digestion was quantified after 1 h of dark exposure applied at different times of the day at 8 dpf in embryos whose lights were turned on at 6:00 or 9:00. The two lighting conditions were compared at each time point (Mann–Whitney U test with Benjamini–Hochberg correction across the three time points). Delaying lights-on to 9:00 significantly suppressed chorion digestion at 12:00–13:00 (*p* = 0.017), whereas at 15:00–16:00 — zeitgeber time (ZT) 6 for the 9:00 group — and at 18:00–19:00, digestion proceeded in both groups to levels that did not differ between the two conditions (*p* = 0.26 and 0.26, respectively). *n* = 10 and 6 (12:00–13:00), 9 and 6 (15:00–16:00), 10 and 6 (18:00–19:00) for the 6:00 and 9:00 lights-on groups, respectively.

We predicted that an extraocular photoreceptor organ controlling hatching exists in the brain, and examined when photoresponsiveness is established. We first examined the maturation of the photoreceptive ability of clownfish embryos. To this end, we exposed 7 dpf and 8 dpf embryos to TRH or ionomycin, a calcium ionophore that activates intracellular calcium signaling, and quantified the extent of chorion digestion under each condition. Chorion digestion was compared among the four treatments at each stage (Kruskal–Wallis test followed by Dunn’s post-hoc test with Benjamini–Hochberg correction, using the DMSO light-treated group as the control; 7 dpf: *p* = 0.0023; 8 dpf: *p* = 4.1 × 10⁻⁹). Dark treatment induced significant chorion digestion only at 8 dpf (Dunn’s test, *p* = 1.6 × 10⁻⁶), whereas no effect was detectable at 7 dpf (*p* = 0.91). By contrast, TRH treatment under light conditions induced chorion digestion at 8 dpf to levels comparable to dark treatment (*p* = 4.3 × 10⁻⁹), and a substantial increase was already apparent at 7 dpf, although this did not reach statistical significance after correction for multiple comparisons (*p* = 0.052; Fig. 3C). Ionomycin significantly increased chorion digestion at 8 dpf (*p* = 5.8 × 10⁻⁴), but its effect was considerably smaller than that of TRH or darkness, and no significant effect was detected at 7 dpf (*p* = 0.91; Fig. 3C). These results indicate that hatching enzyme secretion is not triggered by dark conditions until 8 dpf, but the hatching glands are mature enough to respond to TRH and secrete hatching enzymes as early as 7 dpf. This suggests that the photoresponse or the connection between photoreception and TRH secretion matures at 8 dpf. Next, we investigated when responsiveness to dark conditions was acquired in 8 dpf embryos. We performed hatching experiments by collecting eggs at different times on 8 dpf and quantifying the extent of chorion digestion after incubating for an hour in the dark. At 8 dpf, embryos were incubated with a normal LD cycle (light; 6:00–20:00, dark; 20:00–6:00), and chorion digestion in response to dark incubation occurred 7–13 h (12:00–13:00, 15:00–16:00, and 18:00–19:00) after turning the light on (Fig. 3D). In addition, hatching experiments were performed using irregular LD cycles. When lights were turned on at 6:00, chorion digestion was already underway from 12:00 to 13:00, whereas embryos whose lights were turned on at 9:00 showed essentially no digestion at the same time point, and the two groups were significantly different (Mann–Whitney U test with Benjamini–Hochberg correction, *p* = 0.017). However, between 15:00–16:00 and 18:00–19:00, both groups underwent comparable digestion (*p* = 0.26 and 0.26) (Fig. 3D). Furthermore, hatching occurred when embryos that had been continuously exposed to light throughout the night of day 7 were subjected to 1 h of darkness from 15:00 to 16:00 on day 8 (Fig. S4). In contrast, embryos that had been reared continuously in the dark from day 7 onward did not exhibit dark-induced hatching, even at 15:00–16:00 on day 8. These results indicate that dark-induced hatching is not simply driven by embryos being in darkness during the afternoon of day 8 but rather that the ability to respond to darkness is established by preceding light exposure during the daytime of day 8.

We then performed transcriptome analysis using 8 dpf embryos collected at 10:00, which did not respond to dark conditions, and 8 dpf embryos collected at 16:00, which hatched in response to dark conditions. We predicted that photoreception occurs in the hypothalamus and other deep brain regions, including the hypothalamic TRH neurons, and therefore used RNA extracted from the heads from which the eyes had been surgically removed. A comparison of the morning and afternoon transcriptomes revealed that 51 genes were upregulated and 31 genes were downregulated in the afternoon (Fig. S5A and C), suggesting that only a small number of genes changed their expression around the time when the link between photoperception and hatching was established. Notably, neither the opsin genes nor *trh* was among the 82 differentially expressed genes (Fig. S5D–F), indicating that the photoreceptive and neuroendocrine machinery is already in place in the morning. Instead, the differences were dominated by circadian clock and light-responsive genes (Fig. S5B,C), and RT-qPCR confirmed robust *per1b* and *cry1a* oscillations across 7–8 dpf with peaks in the light phase (Fig. S6), showing that a functional molecular clock operates in the embryonic brain before hatching. Together, these results localize the gating step downstream of photoreceptor gene expression — to signaling, protein-level or circuit-level events rather than to changes in the abundance of the transcripts encoding the pathway.

### Spectral sensitivity analysis

Next, we focused on photoreception upstream of the hatching system in false clownfish. Many light-dependent biological phenomena are characterized by spectral sensitivity, which represents the characteristic response profile to specific wavelengths of light. Spectral sensitivity is important not only for understanding the wavelength dependence of photobiological processes but also for identifying the contributing photoreceptor molecules, as it correlates with the absorption spectra of those molecules. Unlike many light-driven responses, hatching in false clownfish is suppressed by light; therefore, we used very weak light and quantified the degree of hatching suppression as the extent of chorion digestion. Specifically, weak monochromatic light (350, 400, 450, 500, 550, 600, 650, 700, and 750 nm) was generated using a large spectrograph (Okazaki Large Spectrograph, NIBB, Japan; Watanabe et al., 1982) and neutral-density (ND) filters and directed at 8 dpf embryos. The light was applied at three intensity levels (×1, ×3 and ×9), where ×1 is the lowest level, at which the photon flux density was equalized across wavelengths. After 1 h of illumination, egg envelopes were collected and the extent of chorion digestion was quantitatively assessed. At ×9 dim light (9.33 × 10¹¹ photons cm⁻² s⁻¹), chorion digestion was strongly suppressed at 450–550 nm and partially suppressed at 350–400 nm, whereas digestion proceeded at 600–750 nm (Fig. 4A). At ×3 (3.11 × 10¹¹ photons cm⁻² s⁻¹), strong suppression remained confined to 450–550 nm, whereas digestion recovered at 350 nm. At ×1 dim light (1.04 × 10¹¹ photons cm⁻² s⁻¹), however, digestion proceeded fully at most wavelengths, and only illumination at 550 nm still almost completely suppressed chorion digestion (Fig. 4A). Strong suppression was therefore confined to the 450–550 nm range at every intensity tested and contracted to 550 nm as the photon flux density approached threshold. Because the range of effective wavelengths converged on 550 nm at the lowest intensity, the photoreceptor(s) regulating hatching are most sensitive in the green region around 500–550 nm. Teleosts have 30–40 opsin genes, and false clownfish have 34 opsin genes (Fig. S7). As opsin homologues often exhibit similar spectral properties, we examined the clownfish opsin homologues *rhodopsin, exorhodopsin, rh2, vaopsin, opn5L1*, and *parietopsin,* which have been reported to exhibit peak sensitivities at longer wavelengths in zebrafish, medaka, and chickens (Chinen et al., 2003; Matsumoto et al., 2006; Kojima et al., 2008; Tarttelin et al., 2011; Sato et al., 2018; Wada et al., 2018). Red opsin (*lws*), which receives the longest wavelength of light, was excluded because it is not expressed in clownfish embryos (Fig. S8). Cryptochrome, another candidate photoreceptor, was also excluded from the analysis because in vertebrates, it typically has peak sensitivity in the short-wavelength range.

**Fig. 4.**
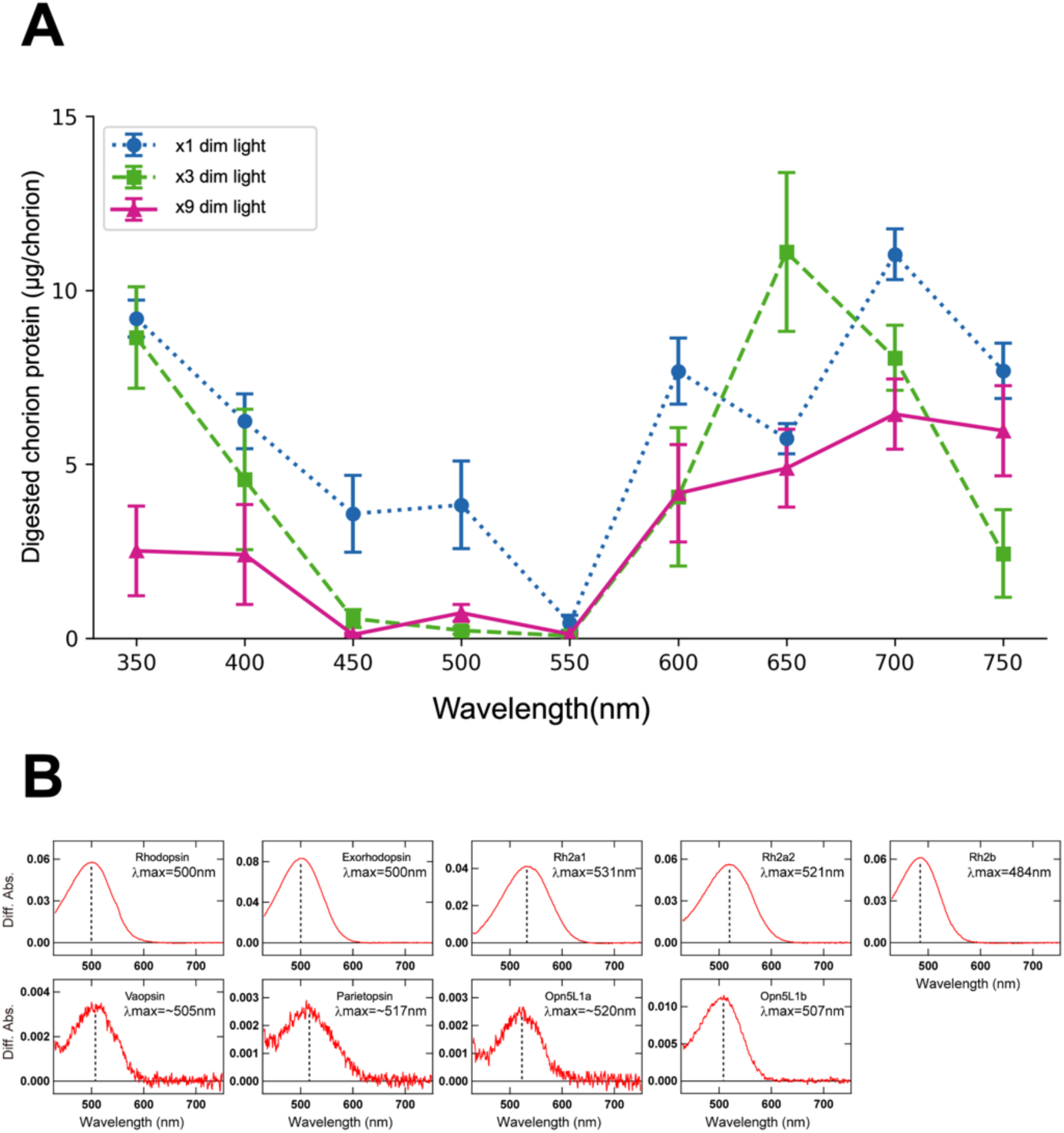
Spectral sensitivity of clownfish hatching and molecular spectrum of clownfish opsins. (A) Wavelength– and intensity-dependent suppression of chorion digestion. Embryos were illuminated for 1 h at each wavelength (n = 3–9 per wavelength and intensity; symbols and error bars show mean ± SEM). At ×9 (9.33 × 10¹¹ photons cm⁻² s⁻¹), chorion digestion was strongly suppressed at 450–550 nm and partially suppressed at 350–400 nm; at ×3 (3.11 × 10¹¹ photons cm⁻² s⁻¹), strong suppression was confined to 450–550 nm. At ×1 (1.04 × 10¹¹ photons cm⁻² s⁻¹), only 550 nm remained suppressive, suggesting that the photoreceptor regulating hatching is most sensitive to green light. (B) Absorption spectra of middle/long-wavelength-sensitive anemonefish opsins. Clownfish homologues of vertebrate opsins reported to be middle/long-wavelength-sensitive were cloned, expressed in HEK293S cells, and their absorption spectra were measured after extraction. All pigments except the cone opsin Rh2b exhibited absorbance maxima (λmax) at or above 500 nm.

*opn5L1, vaopsin*, and *parietopsin* are non-visual opsins whose functions have not yet been fully characterized in teleosts. The molecular absorption maxima (λmax) of Opn5L1a, Opn5L1b, Vaopsin, and Parietopsin are 520 nm, 507 nm, 505 nm, and 517 nm, respectively (Fig. 4B), showing that these opsins have molecular absorbance in the green light range around 500–550 nm consistent with spectral sensitivity. Exorhodopsin is a non-visual opsin generally expressed in the teleost pineal organ and is involved in circadian regulation, and its λmax is 500 nm (Fig. 4B). Rh2 is a visual opsin that is typically expressed in green-sensitive cones. During gene cloning, we identified two distinct *rh2a* sequences: *rh2a1* and *rh2a2*. The λmax values of the three *rh2* homologues—Rh2a1, Rh2a2, and Rh2b—were 531 nm, 521 nm, and 484 nm, respectively (Fig. 4B). Except for Rh2b, these opsins have molecular absorption peaks within the 500–550 nm range identified by spectral sensitivity, positioning them as candidate photoreceptors that regulate hatching. However, the knockout of candidate opsins simply selected based on their molecular spectra and expression data, namely, *exorhodopsin* and *vaopsin* double knockout and *opn5L1a/b/d* triple knockout F0 mosaic crispants, resulted in dark-induced hatching comparable to that of the wild type (Fig. S9).

## Discussion

The timing of fish hatching varies among species, with diverse strategies ranging from those dependent on embryonic developmental stages to those tightly regulated by specific environmental cues, such as light and oxygen levels (Yamagami, 1988; Helvik & Walther, 1992; McAlary & McFarland, 1993; Dimichele & Taylor, 1980; Griem & Martin, 2000). For many years, calcium signaling has been the only known intracellular mechanism to be involved in the regulation of hatching (Schoots et al., 1981); however, TRH and *Trhra* have recently been reported to function as a hatching hormone and its receptor in zebrafish and medaka (Gajbhiye et al., 2024). Yet how environmental cues are integrated into these mechanisms remains largely unknown. In this study, we show that extraocular photoreception is coupled to the conserved TRH–Trhra system, which induces hatching enzyme secretion via Ca²⁺ signaling. *trh* was expressed in the hypothalamic region (Fig. 1A) and exogenous TRH activates intracellular Ca²⁺ signaling via *Trhra* in hatching gland cells, inducing the secretion of hatching enzymes (Fig. 2). The function and expression patterns of *trh* and *trhra* were consistent with those in medaka and zebrafish (Gajbhiye et al., 2024), showing that the TRH–*Trhra*–Ca²⁺– hatching enzyme system is highly conserved and represents a core molecular module regulating hatching across teleosts. We further indicated that, in false clownfish, this conserved downstream mechanism is regulated by upstream extraocular photoreception. Embryos in which the eyes were almost completely absent, generated by CRISPR/Cas9-mediated disruption of *pax6b* and *strip1*, nevertheless digested the chorion in response to darkness and were suppressed by light to the same extent as embryos with intact eyes, with no detectable effect of genotype (Fig. 3A and B). The light information that governs hatching is therefore detected outside the eyes, implying that photoreceptors located elsewhere in the embryo monitor the ambient light environment at the time of hatching. Extraocular photoreception occurs in many vertebrates. Recently, non-visual opsins have been shown to be involved in the regulation of specific light-driven phenomena, including seasonal reproduction and body color changes (García-Fernández et al., 2015; Fukuda et al., 2025). Moreover, even in animals without eyes, various biological phenomena are regulated by light. For example, in jellyfish, light-induced spawning is controlled by *opsin9* expressed in cells that secrete maturation-inducing hormone (MIH) (Quiroga Artigas et al., 2018; Takeda et al., 2018). The central nervous system, including the pineal organ and the hypothalamus, the latter of which harbors TRH neurons, represents a plausible organ for extraocular photoreception. Although the molecular identity of the extraocular photoreceptor(s) controlling hatching remains to be determined, this study establishes a key spectral constraint that should narrow the search.

Spectral sensitivity analysis, which provides crucial information for identifying photoreceptors, revealed that hatching was most effectively suppressed by green light (550 nm) (Fig. 4A). In many light-driven phenomena, spectral sensitivity is known to correlate with the absorption spectra of the underlying photopigments, and this approach for inferring candidate molecules from spectral sensitivity has been widely applied. Because hatching in false clownfish is driven by the dark and suppressed by light, we measured the extent of light suppression using weak monochromatic light. The spectral sensitivity of hatching did not show a similar response across a broad range of wavelengths; rather, a relatively narrow wavelength range (500–550 nm) effectively suppressed hatching, suggesting that a single photoreceptor or a small number of photoreceptors with similar wavelength sensitivities (middle-long wavelength) were involved. Nevertheless, F0 crispants of candidate opsins matching this spectral range, including three *opn5L1* homologues, *vaopsin* and *exorhodopsin*, exhibited dark-induced chorion digestion comparable to that of wild-type embryos (Fig. S9), suggesting the involvement of additional opsins or functional redundancy among the photoreceptors. Identifying precise photoreceptors and delineating the neural circuitry linking them to TRH neurons remains a challenge for future studies. In addition, the fact that hatching is most strongly suppressed by green light also makes ecological sense. As the path length of sunlight in the atmosphere increases at sunset and moonlight is red-shifted compared to daylight, green light constitutes a dominant spectral component of the underwater light environment during twilight before and after sunset in coral reefs (Sweeney et al., 2011). Similarly, twilight around sunset and complete darkness preceding moonrise have been reported as regulatory cues for coral spawning (Lin et al., 2021). Thus, green light sensitivity in hatching is likely adaptive, as it suppresses hatching under relatively bright conditions immediately after sunset or during full moon nights and promotes hatching under complete darkness, where larval dispersal can occur more efficiently.

Our data further revealed that dark responsiveness during hatching is temporally gated rather than constitutively present throughout the day. Delaying the onset of light exposure correspondingly delayed the time at which the embryos became responsive to darkness, indicating that this competence was established relative to the light phase rather than the absolute time of the day (Fig. 3D). Notably, embryos kept in constant darkness from 7 dpf failed to hatch, and dark responsiveness was acquired only after a period of preceding light exposure (Fig. S4), demonstrating that light is not merely permissive but is actively required to establish the capacity to respond to subsequent darkness. Once this competence is established in the afternoon of the hatching day, darkness promotes hatching irrespective of the time at which it is presented; even a dark pulse delayed by an additional day still triggers hatching (Fig. S4), suggesting that embryos were held in a light-suppressed but hatching-competent state rather than losing competence over time. Consistent with the involvement of a circadian mechanism, several clock genes were differentially expressed in the morning and afternoon of hatching (Fig. S5A and C), suggesting that the molecular clock operates in the embryonic brain. However, because our transcriptome analysis sampled only two time points, whether this clock output directly gates the dark responsiveness of hatching or reflects a parallel developmental or diurnal process remains to be determined. Taken together, this study defines several key features of nocturnal hatching in the false clownfish: the conservation of the hatching system, the contribution of extraocular photoreception, the spectral sensitivity, and the temporal gating of dark responsiveness. Nocturnal hatching has been reported in several species of Pomacentridae, including the false clownfish. Therefore, it is possible that the integration of the photoregulatory system with the core TRH–hatching enzyme module at the base of Pomacentridae underlies the acquisition of nocturnal hatching in this lineage, although this remains to be tested phylogenetically. To some extent, this event may have contributed to the adaptive radiation of this lineage in coral reef habitats characterized by high population densities during the daytime.

## Materials and Methods

### Egg incubation

Eggs were isolated from adults and were maintained in a circulating system described in a previous study (Yamanaka et al., 2021) at 27–28°C under a 14/10 LD cycle with lights on at 6:00 AM. Experiments were performed in accordance with the Law for Humane Treatment and Management of Animals of Japan and Kyoto University’s Regulations on Animal Experimentation. The care and use of fish were approved by the Kyoto University Animal Ethics Committee (Approval Code: R4-45 and R5-45).

### In situ HCR

*In situ* HCR was performed according to the manufacturer’s protocol (Nepagene, Japan). The DNA probes were designed against the coding sequences of *trh*, *trhra,* and *lce* (Table S1) and synthesized (Eurofins Genomics, US). The embryos were fixed in 4% paraformaldehyde overnight at 4°C, manually dechorionated, and immersed in 100% methanol at 4°C. After rehydration and prehybridization, the embryos were incubated in hybridization buffer with 20 nM probes overnight at 37°C. Embryos were washed thrice with 0.5× SSCT and incubated with amplification buffer for 5 min at room temperature. The embryos were then incubated with amplification buffer containing a short hairpin amplifier and DAPI for 2 h. After amplification, the embryos were washed several times in PBST and observed under a confocal microscope (LSM710; ZEISS, Germany).

### Chorion digestion assay

The extent of chorion digestion was quantified as the amount of solubilized chorion protein released by hatching enzymes. Embryos at the indicated stages were incubated in 50 mL of seawater in a 200 mL flask at 28 °C for 1 h with shaking at 60 rpm on a Multi Shaker MMS-110 (Tokyo Rikakikai, Japan). Light conditions were provided by room light; for dark conditions, flasks were wrapped in aluminum foil and placed in a light-proof box to exclude light completely. Immediately after the incubation, the egg envelope of each embryo was removed with fine forceps and boiled at 100 °C for 5 min in 2% SDS to extract digested chorion proteins. Each sample consisted of the egg envelope of a single embryo. Protein in the supernatant was quantified with a Pierce™ BCA Protein Assay Kit (Thermo Scientific, US) against a standard curve prepared from the bovine serum albumin (BSA) supplied with the kit, and absorbance was read at 550 nm on a Multiskan FC microplate reader (Thermo Scientific, US).

### Gene knockout

To generate knockouts using the CRISPR/Cas9 system, single guide RNAs (sgRNAs) were designed and synthesized according to the manufacturer’s protocol of CUGA T7 gRNA Synthesis Kit (Nippon Gene, Japan). The target sequences are listed in Table S2. Injection solution including Cas9 protein (250 ng/µL) and sgRNA (100 ng/μL each) for the target genes was injected into the cytosol of fertilized eggs as described in a previous report (Yamanaka et al., 2021).

### Generating GCaMP6s transgenic F0 embryos and calcium imaging

To examine the effect of darkness on Ca²⁺ levels within hatching gland cells, we generated embryos expressing GCaMP6s and mCherry by microinjection as previously described (Yamanaka et al., 2021). On the hatching day, we observed changes in GCaMP6s fluorescence before and 30 min after shading. All fluorescence images were captured at the same exposure time.

### Treatment of TRH/ionomycin

Eggs were incubated for 1 h under light conditions in either 5 μM ionomycin in seawater, 50 μM TRH in seawater, or DMSO in seawater at the same final concentration (vehicle control). The extent of chorion digestion was quantified using the BCA assay as described above. In addition, eggs were incubated for 1 h in the dark in DMSO/seawater, and chorion digestion was quantified in the same manner.

### Dark responsiveness under different LD cycles

To examine whether the duration of the light phase immediately preceding hatching affected dark responsiveness, embryos were reared under an LD 14:10 cycle (lights on at 6:00, lights off at 20:00) until day 7. On day 8, lights were turned on at either 6:00 or 9:00, and eggs were exposed to 1 h of darkness, 3–12 h after lights were turned on. The extent of chorion digestion was quantified using the method described above.

### Spectral sensitivity analysis

Spectral sensitivities were analyzed using monochromatic light within the range of 350– 750 nm using the Okazaki Large Spectrograph at the National Institute for Basic Biology (Watanabe et al., 1982). The irradiance at the specimen position was measured using a photon meter (QTM-101, Monotech, Japan) and converted to photon flux density (PFD). The ND filters were used to generate three intensity levels at each wavelength (nominal 1-, 3-, and 9-fold). Measurements were made at the lowest level, at which the PFD was equalized across wavelengths (mean 1.72 nmol m⁻² s⁻¹, equivalent to 1.04 × 10¹¹ photons cm⁻² s⁻¹); the corresponding means for the 3– and 9-fold levels were 5.16 and 15.5 nmol m⁻² s⁻¹ (3.11 × 10¹¹ and 9.33 × 10¹¹ photons cm⁻² s⁻¹). The wavelength– and level-specific values are listed in Table S3. We quantified the extent of chorion digestion in response to monochromatic light. On the afternoon of the hatching day, eggs were placed in 50 mL of seawater in a 200 mL flask and exposed to monochromatic light (350, 400, 450, 500, 550, 600, 650, 700, and 750 nm) while incubating them on a handmade small shaker for 60 min at 28°C. After incubation, EDTA was added to the seawater to terminate the chorion digestion reaction and egg envelopes were collected using fine tweezers. The extent of chorion digestion was quantified as described previously.

### Spectrum analysis of opsins

The coding sequences of opsins (*rhodopsin, exorhodopsin, rh2a1, rh2a2, rh2b*, *vaopsin* and *parietopsin*) with the epitope sequence of the anti-bovine rhodopsin monoclonal antibody Rho1D4 (ETSQVAPA) at the C-terminus were introduced into the mammalian expression vector pCAGGS. To improve the expression levels of opsin proteins in cultured cells, the N– and C-termini of the coding sequences of *opn5L1a* and *opn5L1b* were replaced with those of *Xenopus tropicalis opn5m* according to a previous paper (Sato et al., 2018). The modified sequences of *opn5L1a* and *opn5L1b* were tagged with the epitope sequence of Rho1D4 at the C-terminus, followed by the introduction into the pCAGGS vector. The plasmid DNA was transfected into HEK293S cells using the calcium phosphate method. 5 μM all-trans retinal (*opn5L1a* and *opn5L1b*) or 11-cis retinal (other opsins) was added to the medium 24 hr after transfection and the cells were kept in the dark until they were collected 48 hr after transfection. The reconstituted photo-pigments were extracted from cell membranes using 1% dodecyl maltoside (DDM) in Buffer A (50 mM HEPES, 140 mM NaCl, pH 6.5). The absorption spectra of the solubilized opsins were recorded before and after light irradiation using a spectrophotometer (UV2600, Shimadzu, Japan) and an optical cell (width, 2 mm; light path, 1 cm). The extracts from cell membranes in the presence of 5 mM NH₂OH (*rh2a1, rh2a2* and *rh2b*) or 50 mM NH₂OH (*rhodopsin, exorhodopsin, vaopsin* and *parietopsin*) or in the absence of NH₂OH (*opn5L1a* and *opn5L1b*) were irradiated with orange light (> 560 nm) which was generated by a halogen lamp (Master HILUX-HR, Rikagaku Seiki, Japan) and passed through an O-58 cutoff filter (Toshiba, Japan). The absorption spectra of opsins were calculated by subtracting the spectrum measured after irradiation from that measured before irradiation.

## Statistical analysis

Data are shown as individual data points with boxplots (median, interquartile range, and 1.5× IQR whiskers). Two-group comparisons were analyzed using the Mann–Whitney U test, and multi-group comparisons were performed using the Kruskal–Wallis test, followed by Dunn’s post-hoc test (Benjamini–Hochberg correction). Statistical significance was set at *p* < 0.05. Two-factor comparisons (genotype × light conditions) were performed using the Scheirer–Ray–Hare test. Proportions were compared with Fisher’s exact test or the χ² test, and group means of transcript abundance were compared with Welch’s t-test, in both cases with Benjamini–Hochberg correction. The analyses used Python 3.10 (SciPy, scikit-posthocs).

## Acknowledgements

We thank Chiyo Takagi, Kentaro Hayashi and Rie Hara for the management of the Okazaki Large Spectrograph facility and for their technical assistance with the spectral sensitivity experiments.

## Author Contributions

Conceptualization, S.Y. and M.K.; Methodology, S.Y., M.K., T.Yamashita., and Y.K.; Investigation, S.Y., T.Yamashita., T.Yamanaka.; Resources, Y.K. and H.T.; Writing – Original Draft, S.Y.; Writing-Review & Editing, all authors; Funding Acquisition, S.Y., M.K., and T.Yamashita.

## Data Availability

The RNA-seq reads data have been deposited in the DDBJ Sequence Read Archive (DRA) under BioProject accession PRJDB45661 (BioSample accessions SAMD02024710–SAMD02024715; Run accessions DRR1092647–DRR1092652).

## Supplementary Information

### Methods

#### Evaluation of mutation rate in injected embryos

After hatching experiments, HMA was performed to investigate the extent of CRISPR/Cas9-induced mutations. A short fragment < 200 bp containing the target site of the sgRNA was amplified by PCR using a primer set (Table S4). The amplicons were analyzed using a microchip electrophoresis system (MCE-202 MultiNA; Shimadzu, Japan) with a DNA-500 reagent kit. In addition, for *pax6b* and *strip1*, the same genomic DNA was used as a template for PCR amplification with individually indexed primers, and the products were pooled and subjected to amplicon sequencing on an Illumina MiSeq platform to quantify the variety and relative abundance of mutant sequences in each individual (Bioengineering Lab, Japan).

### Clock gene RT-qPCR

Eggs were reared under a normal LD cycle (LD 14:10; lights on at 6:00, lights off at 20:00), and four tubes of 10 embryos each were sampled every 4 h from 00:00 on day 7 to 20:00 on day 8. Total RNA was extracted using ISOGEN (Nippon Gene, Japan), reverse-transcribed with SuperScript III (Invitrogen, USA), and amplified with THUNDERBIRD qPCR Mix (TOYOBO, Japan). Expression levels were quantified by the ΔΔCt method, using beta-actin (*actnb*) as the reference gene. The primers used are listed in Table S4.

### Transcriptome analysis

RNA was extracted from head region, excluding eyes and lower jaw at 10:00 and 16:00 on the 8th day using ISOGEN with a spin column kit (Nippon Gene, Japan) according to the manufacturer’s protocol. Then, RNA sequencing was performed by Rhelixa, Inc. (Japan) using NovaSeq (Illumina, USA). Sequence data were trimmed by fastp (Chen et al., 2018) and then subjected to mapping and read count by STAR (Dobin et al., 2013) and RSEM (Li and Dewey, 2011). Differentially expressed genes (DEGs) between ZT4 and ZT10 were identified using edgeR (Robinson et al., 2010). Gene IDs were subsequently converted to zebrafish orthologs using biomaRt (Durinck et al., 2009), and Gene Ontology (GO) enrichment analysis was performed.

### Phylogenetic analysis of opsins

Opsin homologous genes of zebrafish, medaka, and false clownfish were obtained from publicly available sequences in the NCBI database. Multiple amino acid sequences were aligned using MAFFT program (Katoh & Standley, 2013) and trimmed using trimAl (Capella-Gutiérrez et al., 2009). Phylogenetic trees were constructed using the maximum likelihood method with the WAG (Whelan & Goldman, 2001) + Gamma model and bootstrap analysis (1,000 replicates) under the PROTGAMMAAUTO option to determine the best-scoring topology using RAxML (Stamatakis, 2014).

---

**Fig. S1.**
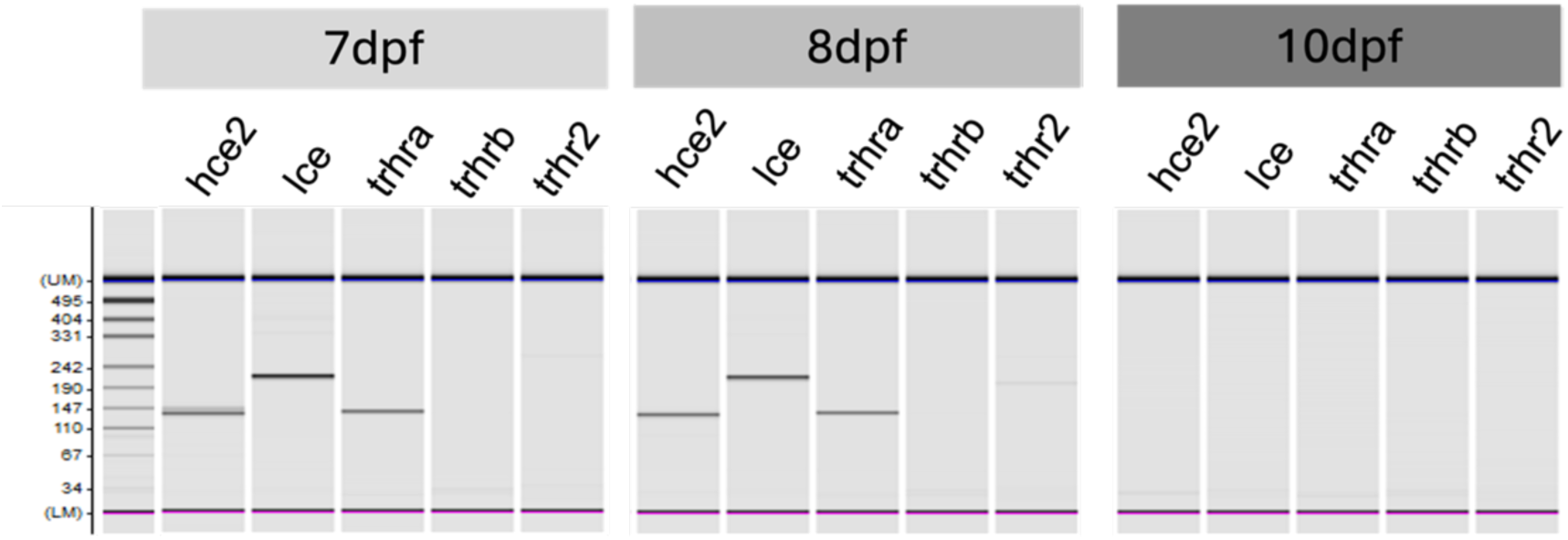
Expression of TRH receptors and hatching enzymes in the trunk region where hatching glands are located. According to RT-PCR, at the pre-hatching stage *trhra* and hatching enzymes are expressed; however, their expression is lost after hatching, although *trhrb* and *trhr2* were not detected at any stage.

**Fig. S2.**
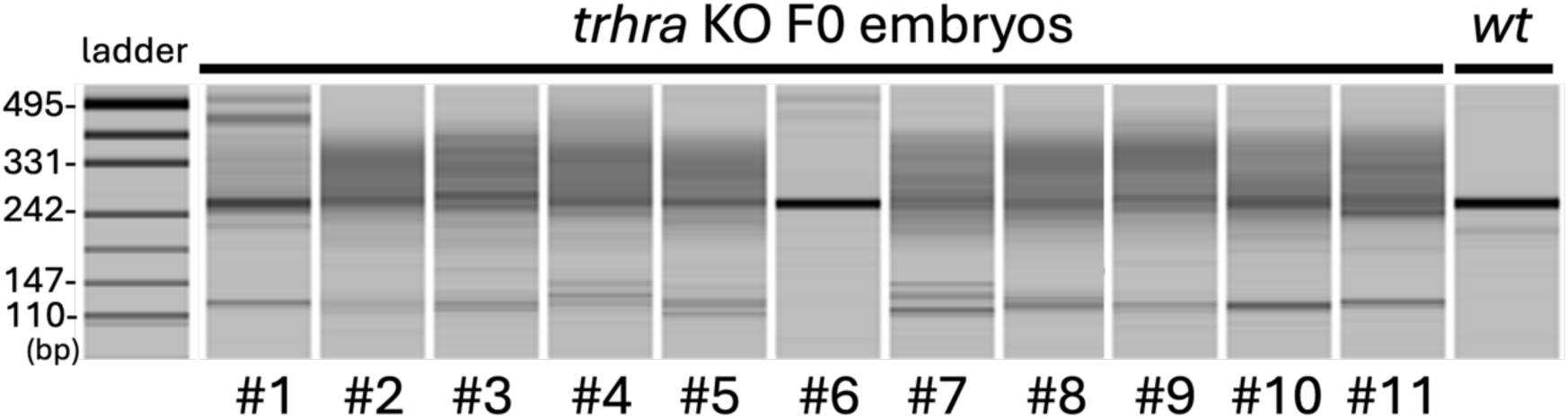
Heteroduplex mobility assay of *trhra* F0 crispants. Almost all injected embryos showed multiple bands in the PCR products including the CRISPR target site at *trhra* indicating that *trhra* was effectively mutagenized.

**Fig. S3.**
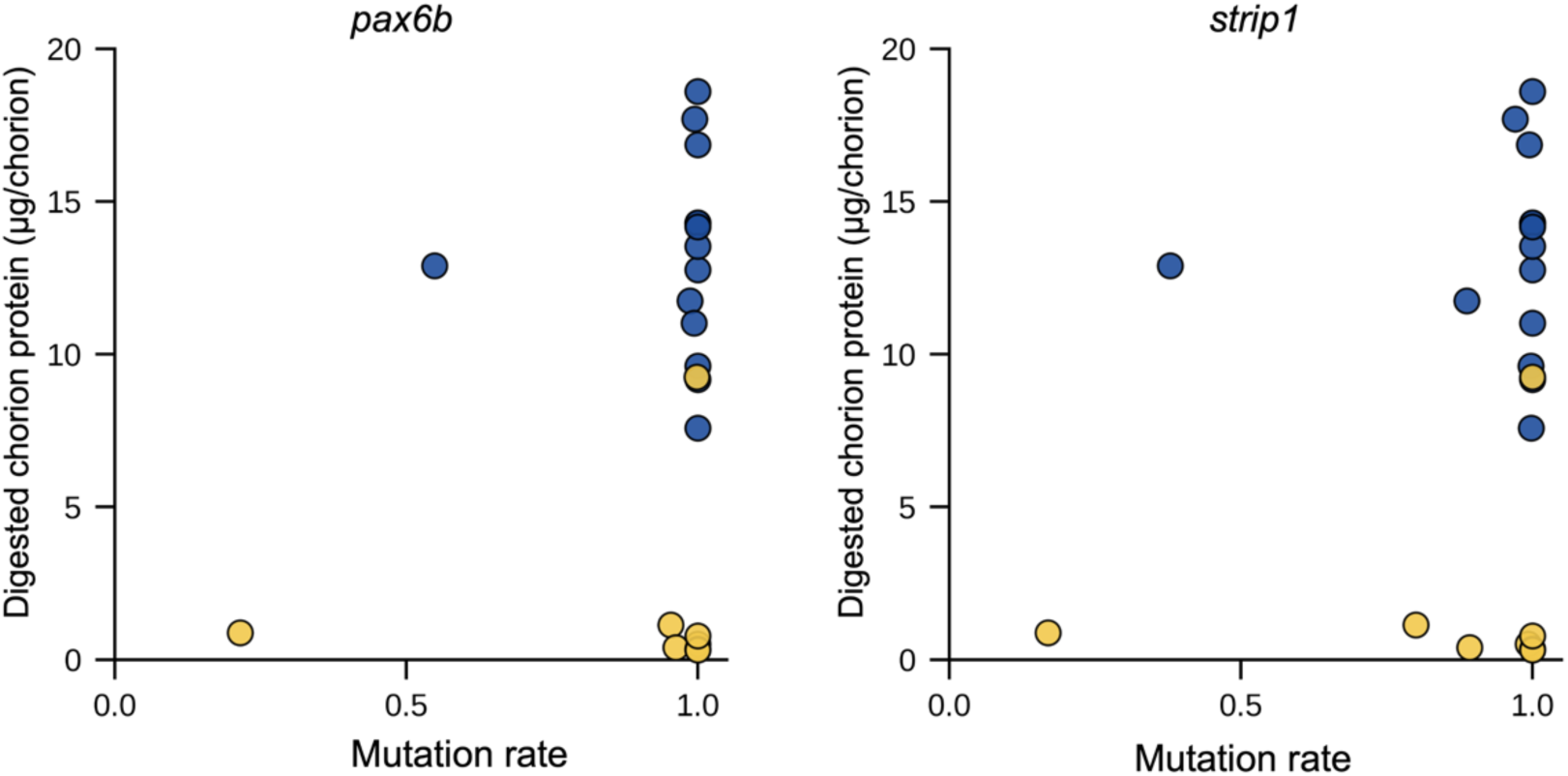
Dot plot of mutation rate at eye-development gene loci against chorion digestion. Individually labeled embryos were exposed to light or darkness for 1 h. Chorion digestion was quantified from the egg envelope of each embryo, and genomic DNA was extracted from the corresponding whole embryo. Regions including the CRISPR target sites in *pax6b* and *strip1* were amplified by PCR from each individual genome, and the proportion of mutated reads was determined by amplicon sequencing. Chorion digestion did not correlate with mutation rate at either locus under either condition (Spearman’s rank correlation: *pax6b*, ρ = 0.09, p = 0.77 in the dark and ρ =−0.58, p = 0.13 in the light; *strip1*, ρ = 0.05, p = 0.88 in the dark and ρ = −0.33, p = 0.43 in the light; *n* = 14 and 8 for dark and light, respectively).

**Fig. S4.**
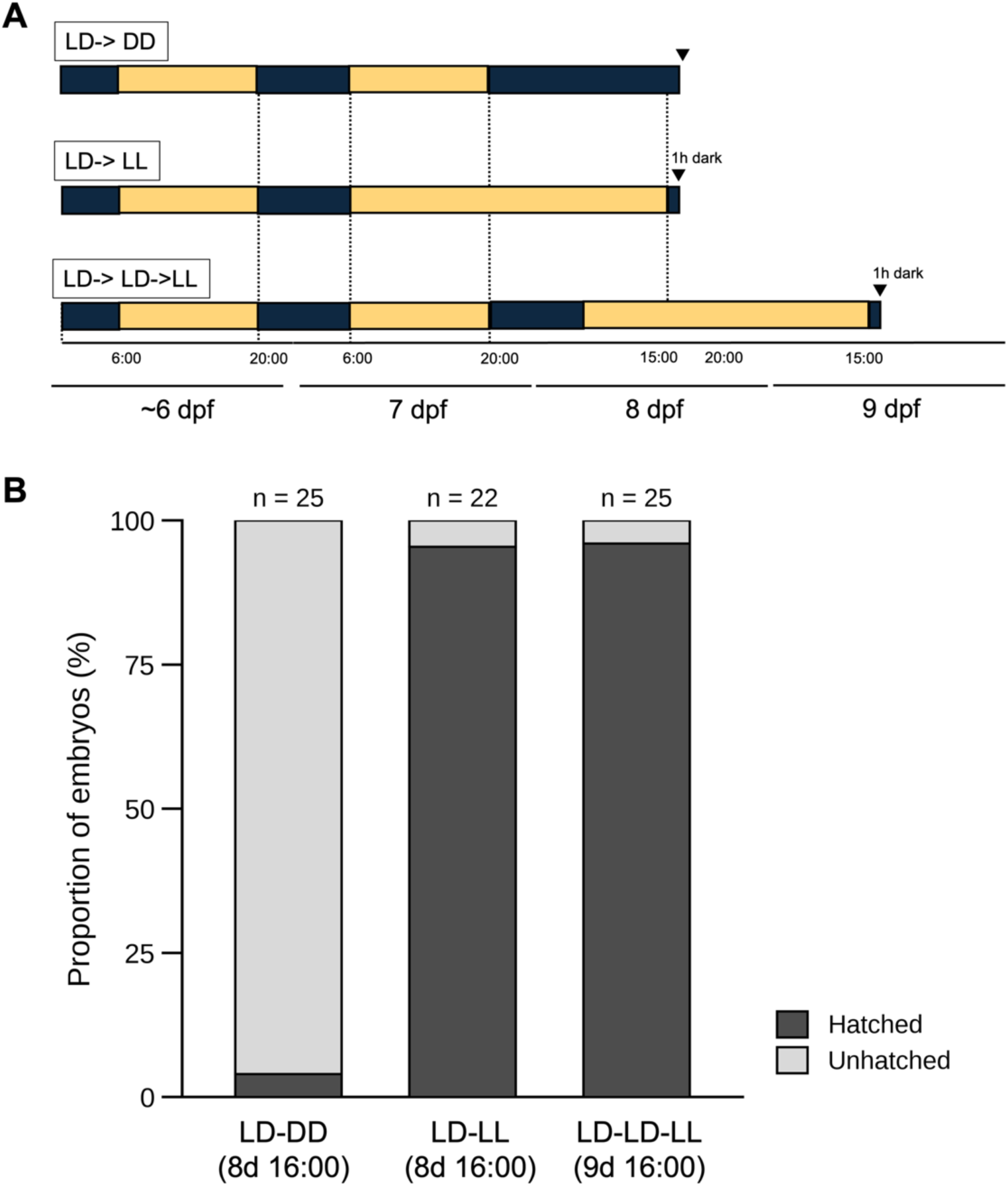
Hatching requires a preceding light phase in addition to a dark transition, and is suppressed by light at any time from the afternoon of 8 dpf. (A) Light regimes. Embryos were reared under LD 14:10 (lights on 6:00) until 7 dpf, then transferred to constant darkness (LD–DD), or to constant light with a 1 h dark pulse at 15:00–16:00 on 8 dpf (LD–LL) or on 9 dpf after one additional LD cycle (LD– LD–LL). (B) Proportion of embryos hatched, scored at 16:00 immediately after the dark pulse. *n*, number of embryos. Hatching failed under constant darkness (1/25) but occurred in almost all embryos given a dark pulse after a light phase (21/22 and 24/25; *p < 0.0001 vs LD–DD*, Fisher’s exact test). Darkness at the appropriate time of day is therefore not by itself sufficient to trigger hatching: preceding light exposure is also required. Furthermore, delaying the dark pulse by 24 h did not reduce hatching, indicating that embryos remain competent but are actively held by light.

**Fig. S5.**
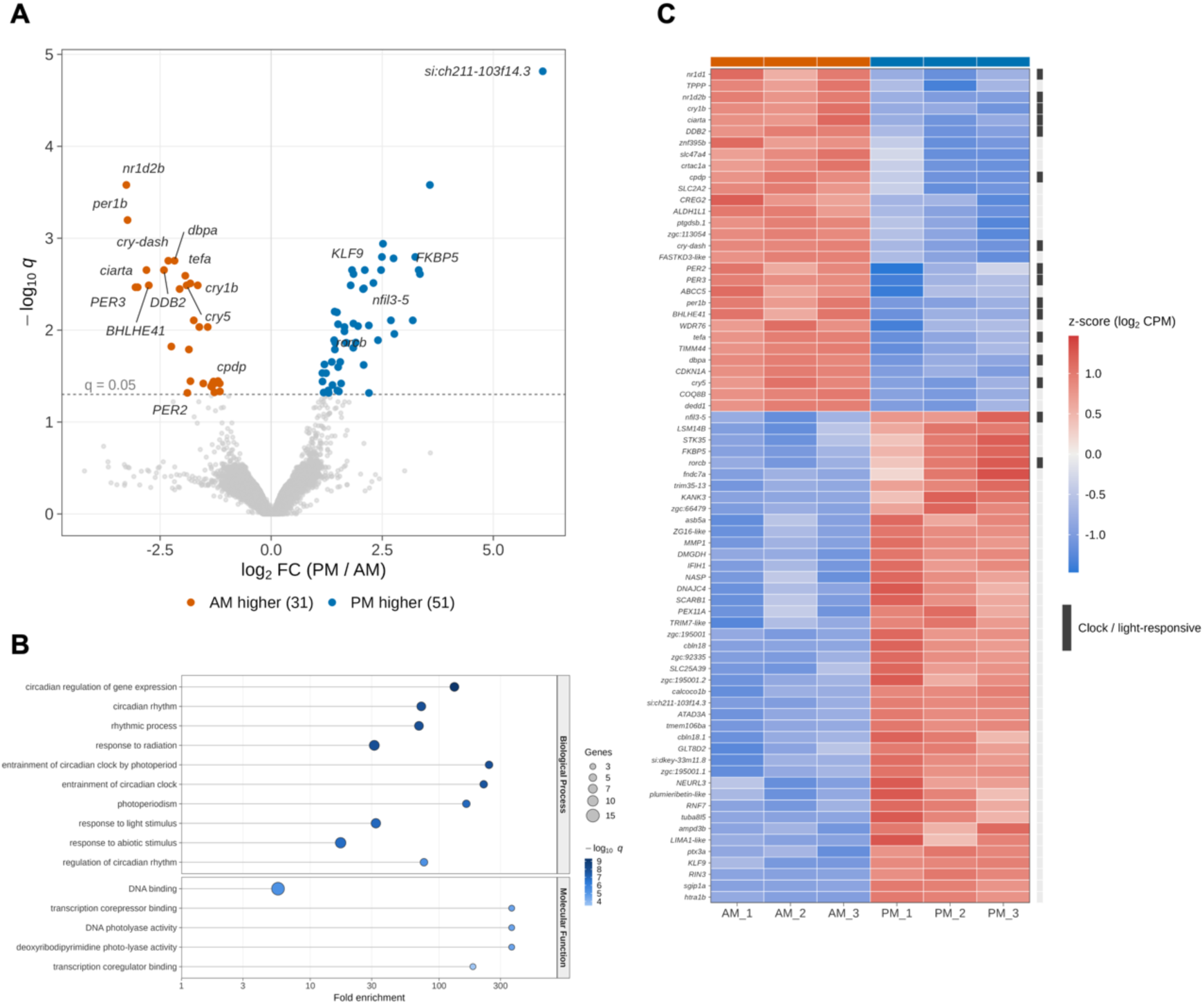
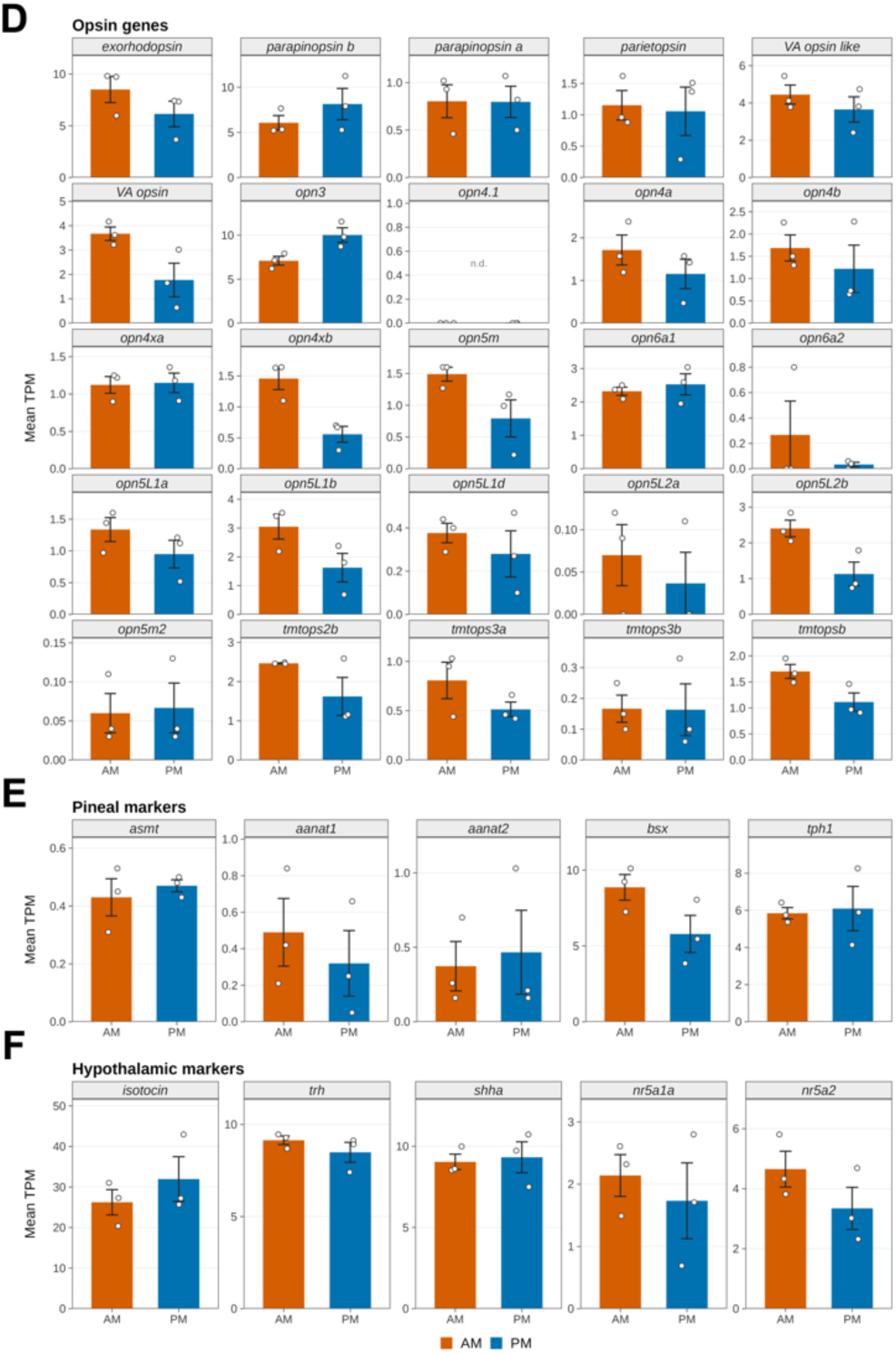
Hatching-day brain transcriptome differs between morning and afternoon in circadian clock and light-responsive genes. (A) Volcano plot of gene expression differences between the morning (AM) and afternoon (PM) of hatching day (brain region, *n* = 3 per group). The log₂ fold change (PM/AM) was calculated from group means of TMM-normalized CPM, and *q* values are Benjamini–Hochberg-adjusted *p* values from edgeR. Genes with *q* < 0.05 are colored (orange, higher in AM, *n* = 31; blue, higher in PM, *n* = 51); the dashed line marks *q* = 0.05. Circadian clock and light-responsive genes and the most strongly PM-biased genes are labeled. (B) Gene Ontology terms over-represented among AM-biased differentially expressed genes (Fisher’s exact test with Benjamini–Hochberg false discovery rate (FDR), using zebrafish GO annotation transferred via orthology). Dot position indicates fold enrichment (log scale), dot size the number of genes annotated to the term, and color −log₁₀ q. The top 15 terms are shown. No term was significantly enriched among PM-biased genes. (C) Heatmap of the 73 protein-coding differentially expressed genes (*q* < 0.05). Color shows the per-gene z-score of TMM-normalized log₂ CPM; rows are ordered by hierarchical clustering. The grey bar on the right marks circadian clock and light-responsive genes. (D–F) Expression (mean TPM) of (D) opsin genes, (E) pineal markers and (F) hypothalamic markers. Bars, group mean (*n* = 3); error bars, standard error of the mean (SEM); open circles, individual replicates (AM, orange; PM, blue). The y axis is scaled independently for each gene. Opsins were expressed at generally low levels (median 1.14 TPM), the exceptions being *opn3*, *exorhodopsin* and *parapinopsinb* (7.1–8.6 TPM); *opn4.1* was not detected (n.d.). Among the pineal markers, *bsx* and *tph1* were clearly expressed (6.0–7.3 TPM), whereas the melatonin-synthesis enzymes *asmt*, *aanat1* and *aanat2* were all close to the detection limit (0.40–0.45 TPM). All hypothalamic markers were clearly expressed (1.9–29.1 TPM). No gene in D–F differed significantly between AM and PM (Welch’s *t*-test with Benjamini–Hochberg FDR, all *q* > 0.50).

**Fig. S6.**
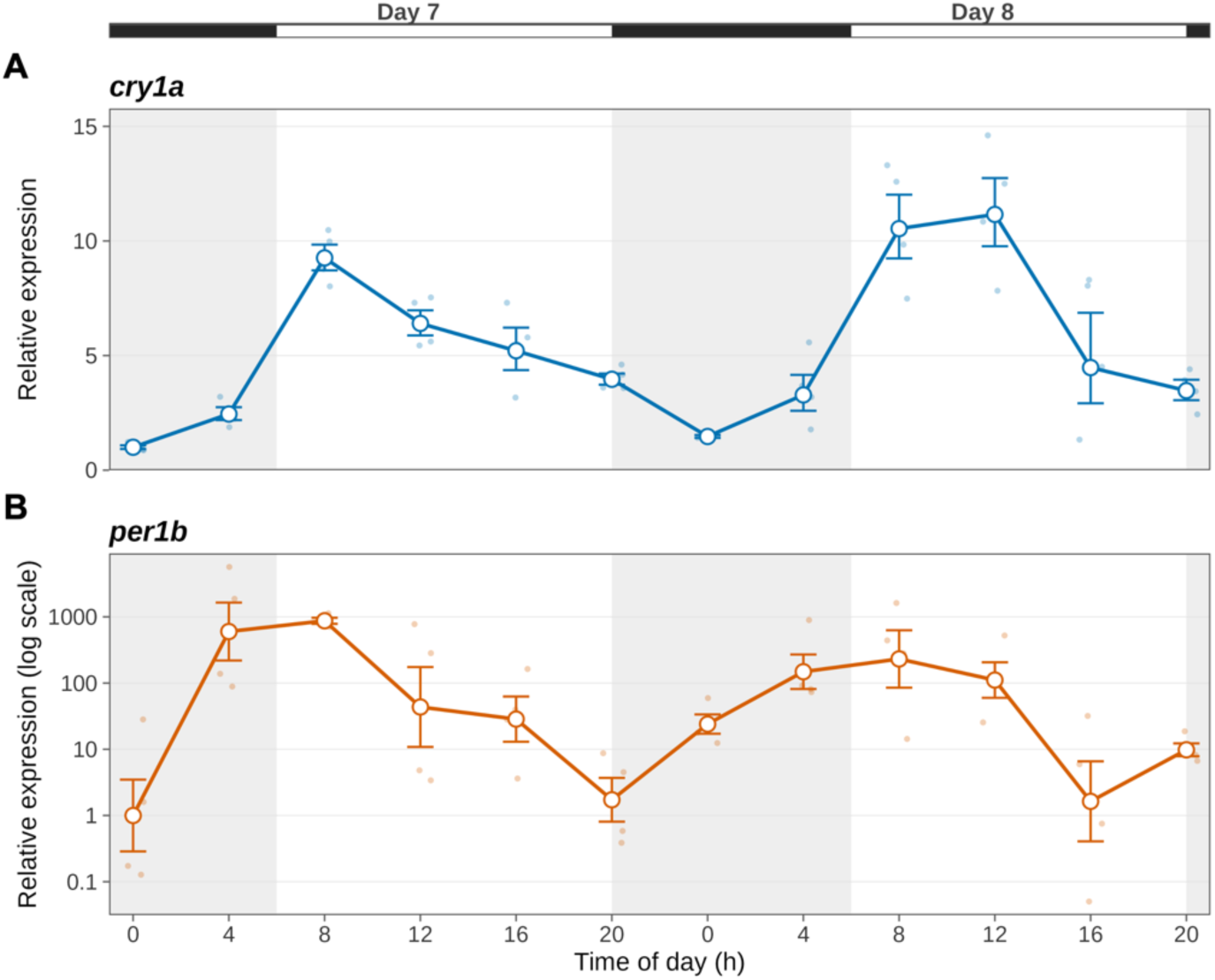
Diel expression of *cry1a* and *per1b* at late stages. RT-qPCR of (a) *cry1a* and (b) *per1b* in whole embryos, sampled every 4 h from 00:00 on day 7 to 20:00 on day 8 (dpf). Levels were normalized to *actnb* and are expressed relative to the lowest time point (day 7, 00:00 = 1; 2^−ΔΔCt). Open circles, mean of four biological replicates, each assayed in technical duplicate; error bars, ± SEM computed on ΔCt and back-transformed (hence asymmetric); small filled circles, individual replicates. Note the logarithmic ordinate in (b). The bars above and shading within the panels denote the light–dark cycle (LD 14:10, lights on at 06:00; ZT0 = 06:00). For *per1b*, 3–5 of the 8 wells gave no Cq at the three trough time points and were assigned Cq = 40; fold changes there were conservative estimates.

**Fig. S7.**
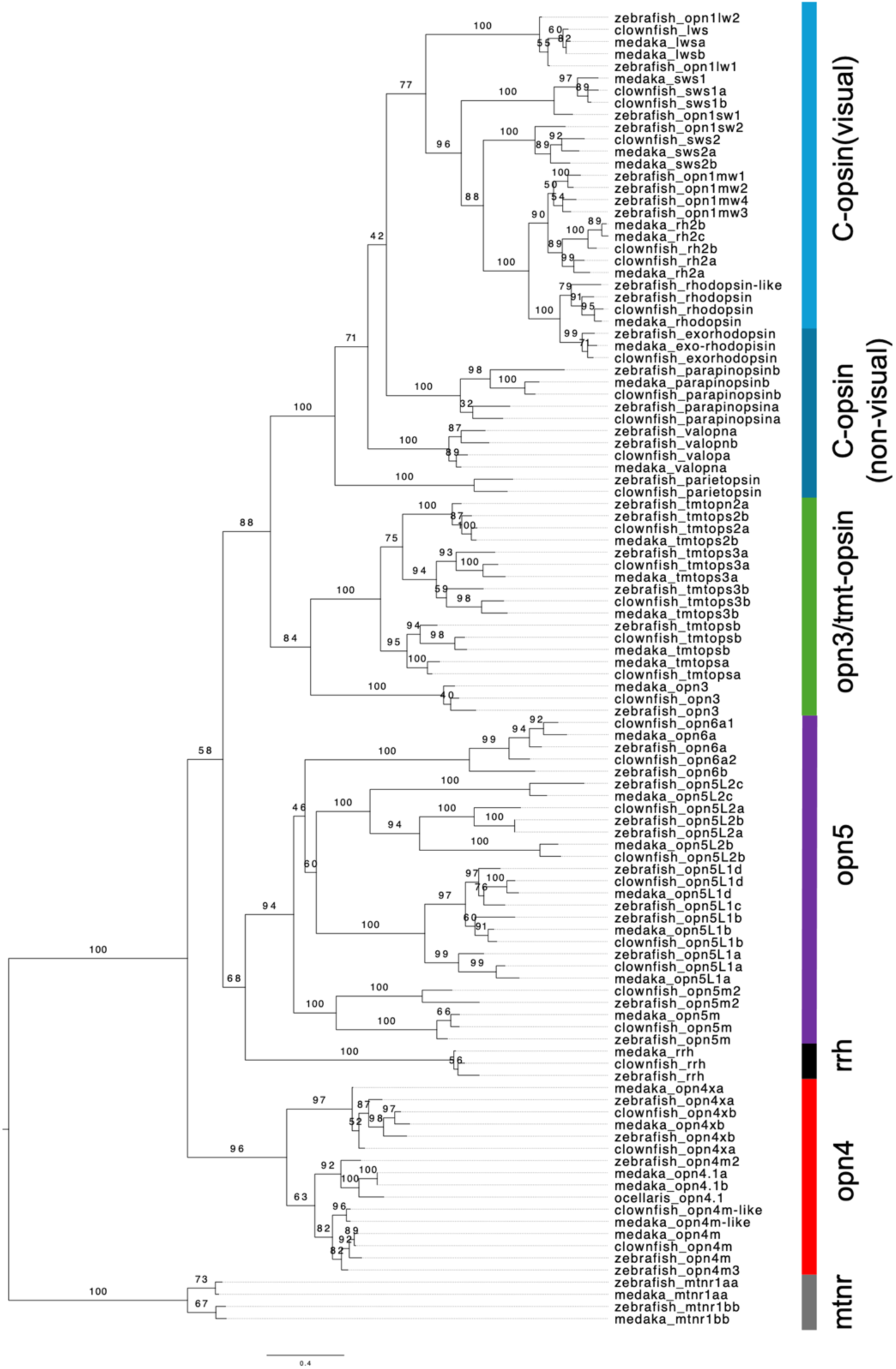
Phylogenetic tree of opsin genes of false clownfish, zebrafish and medaka. False clownfish have 34 opsin genes basically conserved in teleost.

**Fig. S8.**
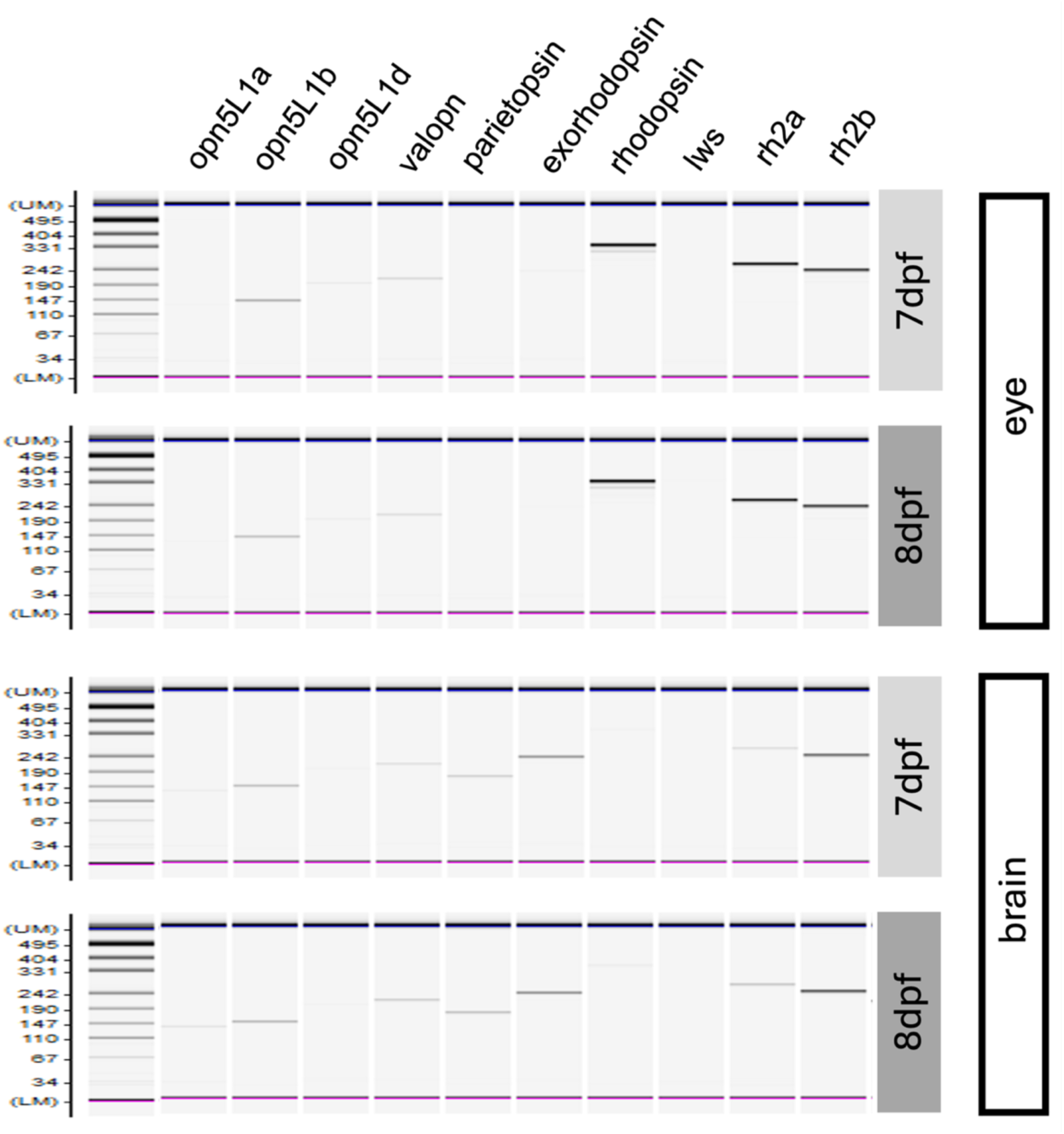
Expression of middle/long-wavelength-sensitive opsin genes of clownfish in eye and brain of embryos. RT-PCR analysis of ten middle/long-wavelength-sensitive opsin genes in the eye and brain of 7 and 8 dpf embryos. Most opsins were detected in both the eye and brain at both stages, with tissue-specific differences in relative band intensity (e.g., *exorhodopsin* and *rh2* paralogs were prominent in the eye, whereas *parietopsin* was more evident in the brain). Amplicons for *lws* and *opn5L1d* were absent in all four conditions, indicating that these opsins were not expressed at detectable levels in pre-hatching embryos.

**Fig. S9.**
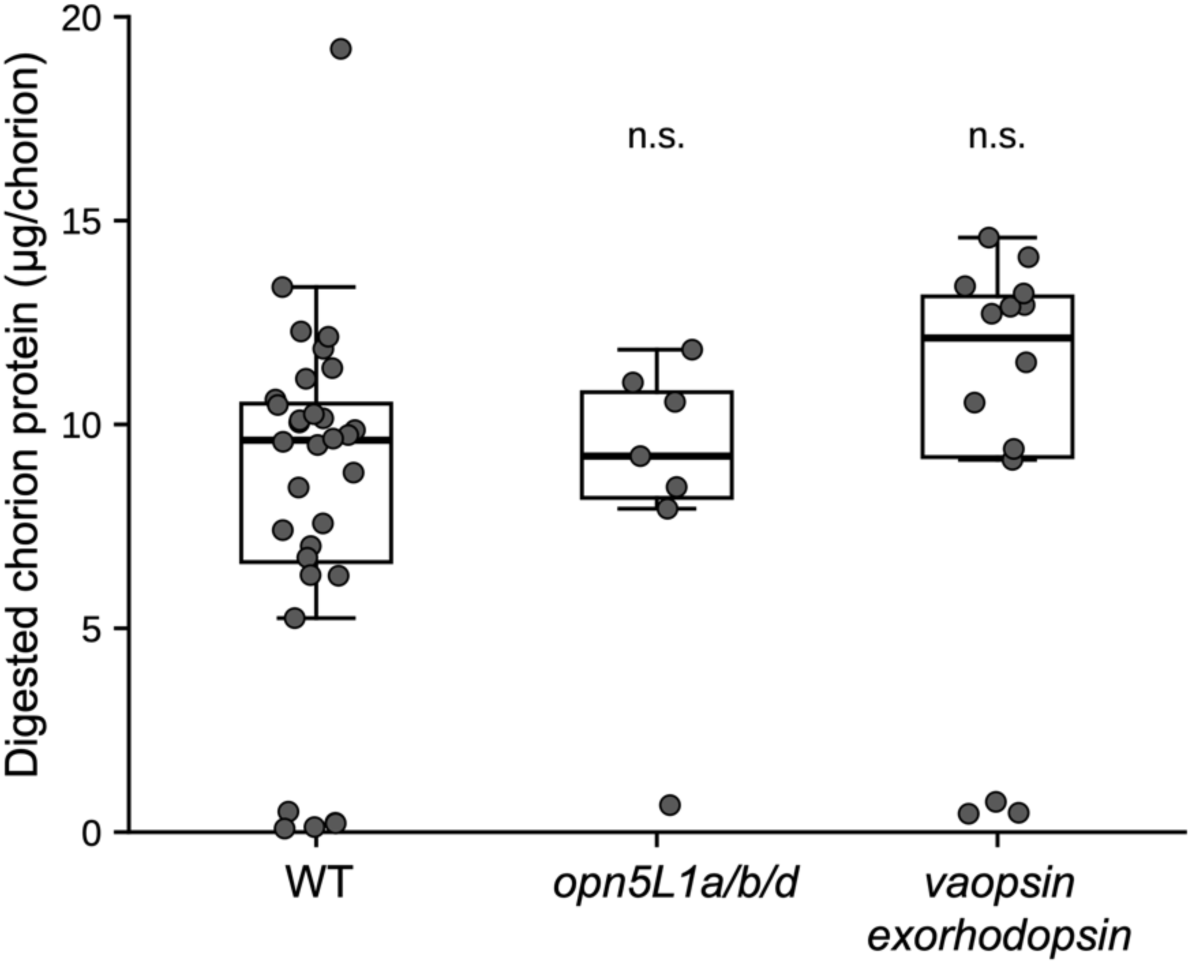
Chorion digestion in mosaic knockout embryos of non-visual opsins. F0 mosaic knockouts were generated for middle/long-wavelength-sensitive non-visual opsins (*opn5L1* homologues, *vaopsin*, and *exorhodopsin*). None of the knockout embryos showed premature hatching or impaired dark-induced hatching at 8 dpf, and chorion digestion did not differ significantly from that of same-day wild-type controls (Kruskal–Wallis test, *p* = 0.13; Dunn’s test with Benjamini–Hochberg correction, *p* = 0.83 for *opn5L1a/b/d* and *p* = 0.14 for *vaopsin*/*exorhodopsin*). *n* = 32 (wild type), 7 (*opn5L1a/b/d*), and 14 (*exorhodopsin*/*vaopsin*).

**Table S1.**
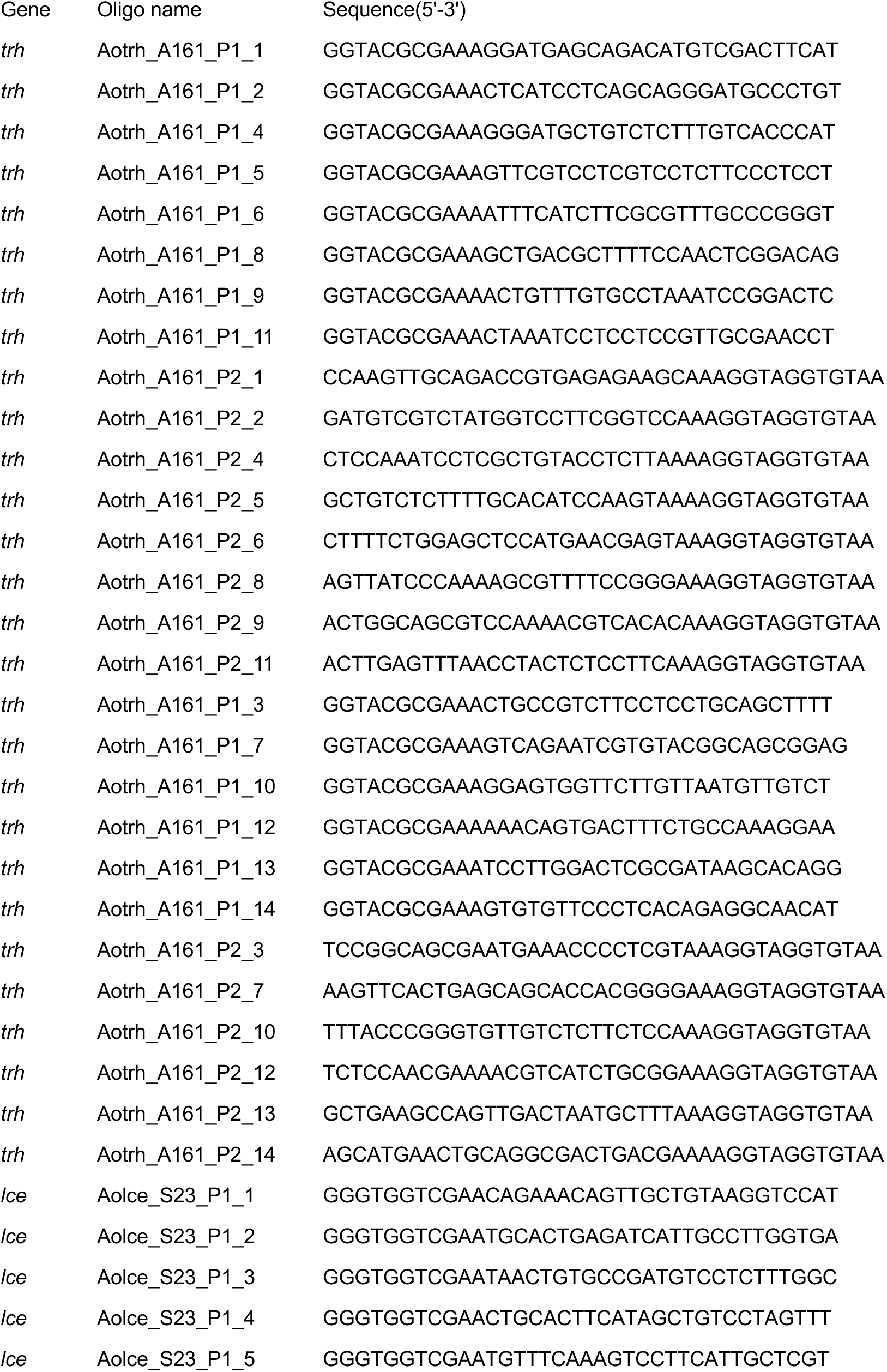
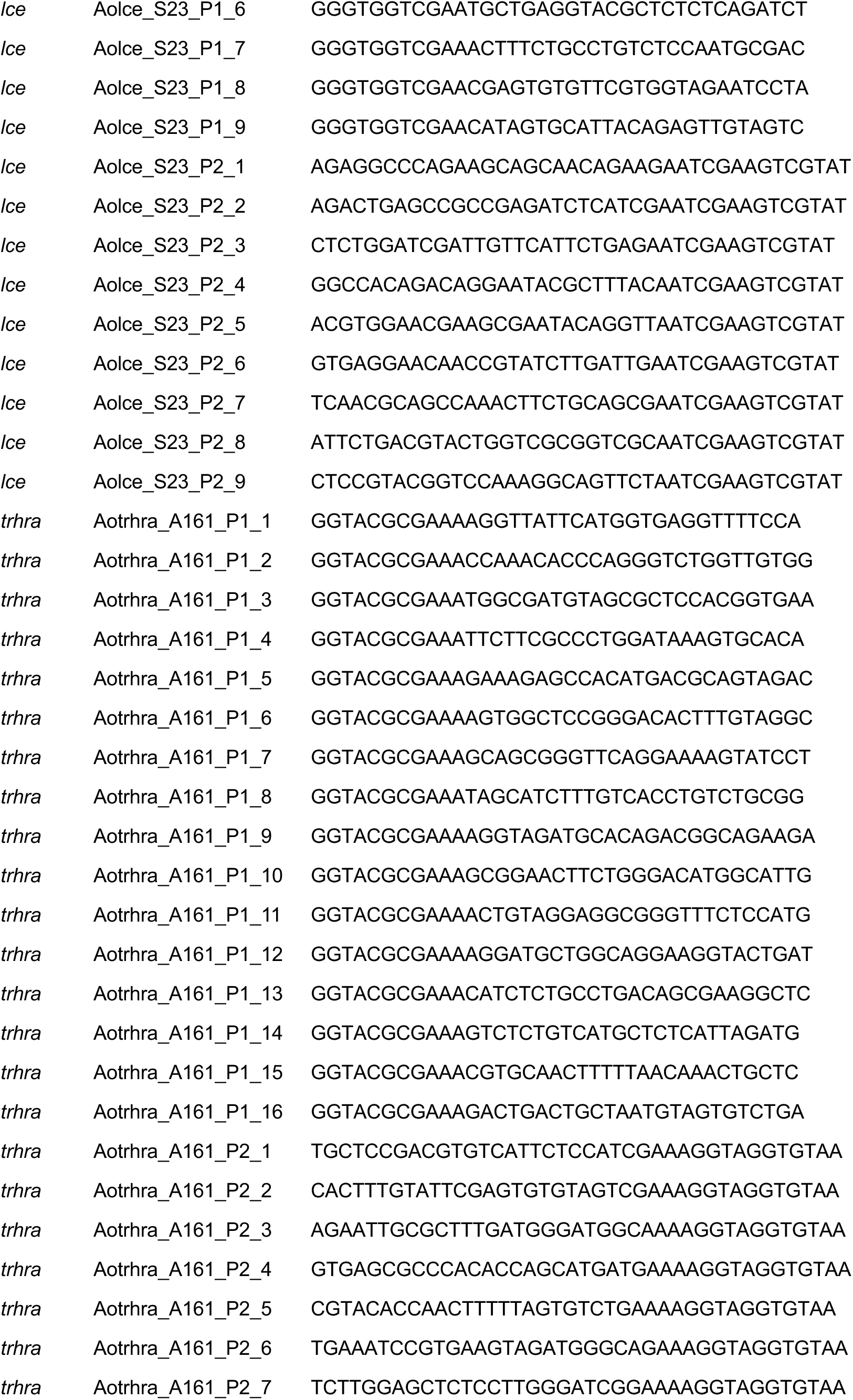
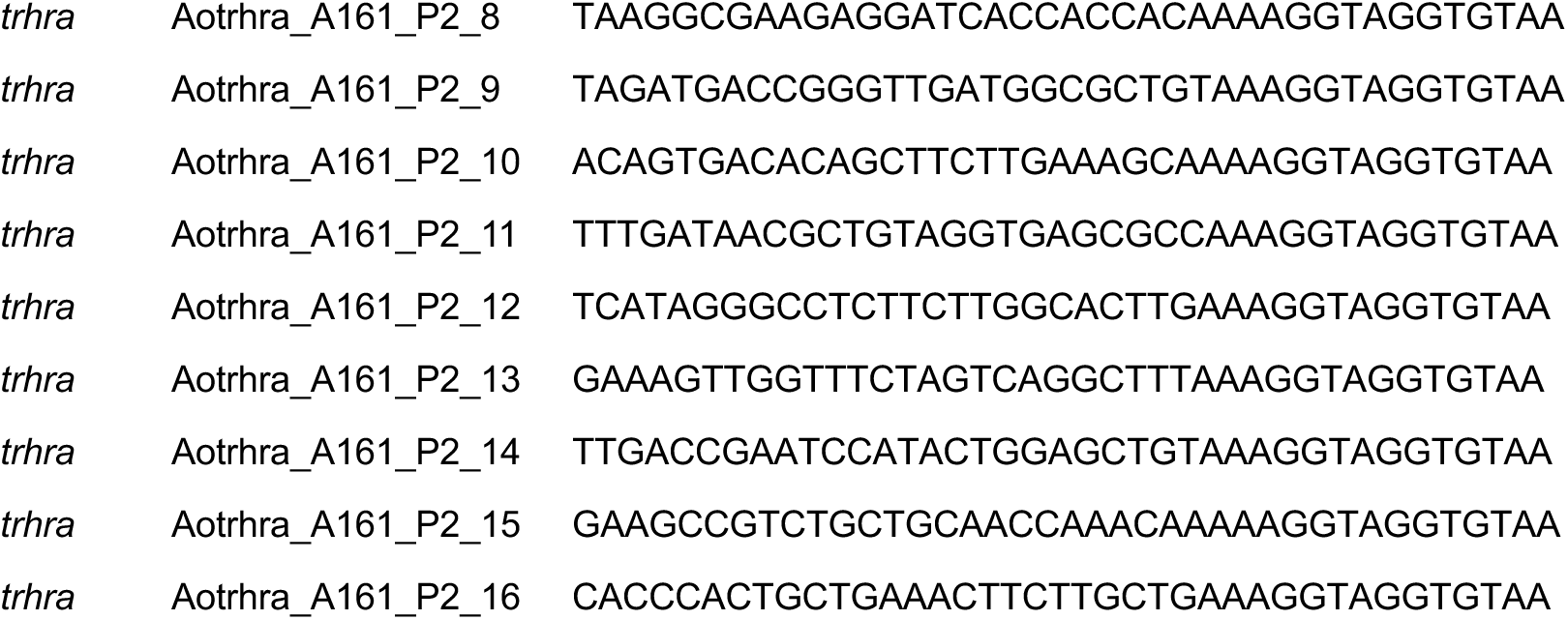
Oligonucleotide probe sets used for *in situ* HCR detection of *trh*, *lce* (hatching enzyme) and *trhra*.

**Table S2.**
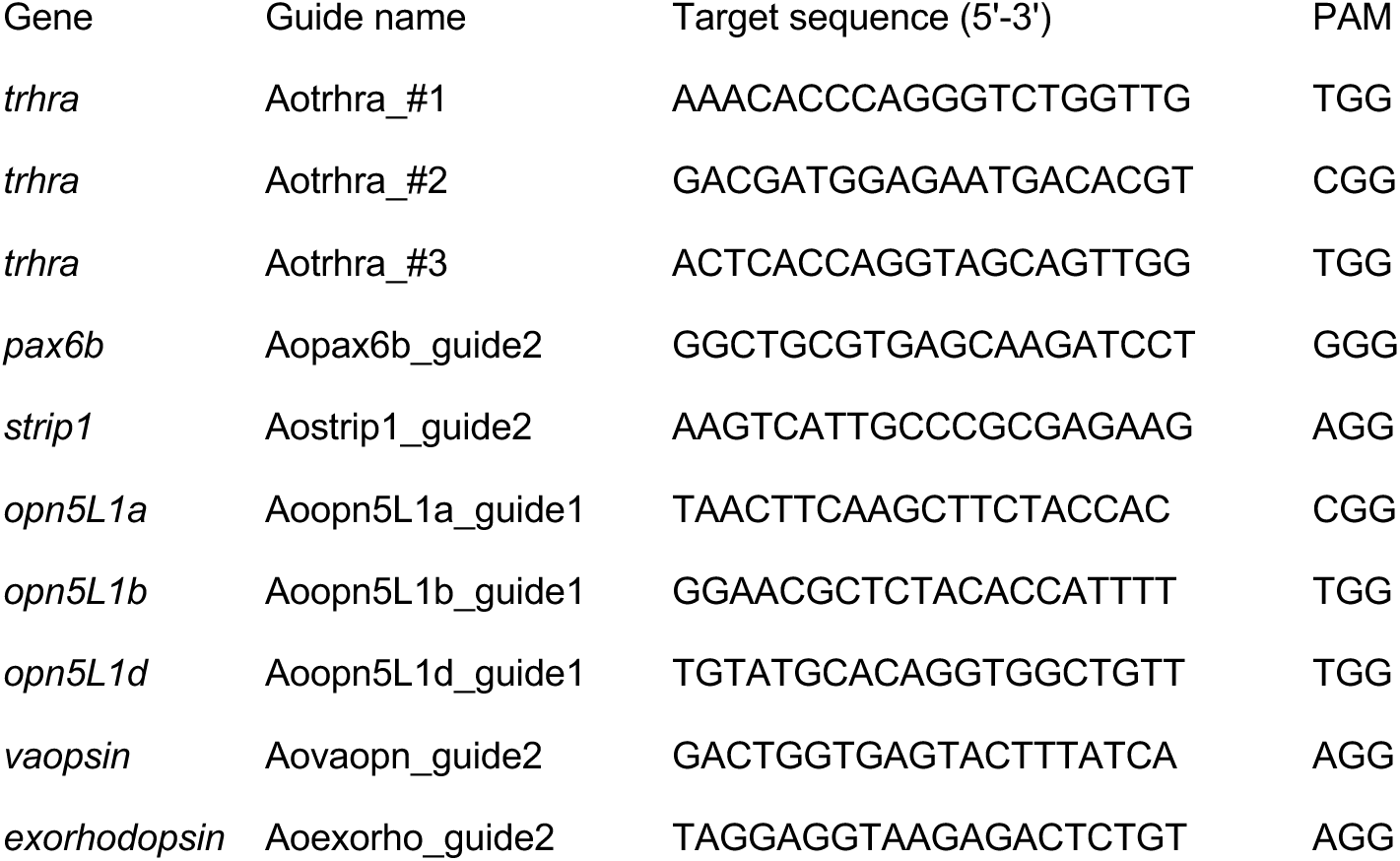
Guide RNA target sequences and adjacent PAM sequences used to generate knockout embryos for *trhra*, *pax6b*, *strip1* and opsin genes.

**Table S3.** Spectral irradiance and photon flux density of the monochromatic light used in this study. Irradiance was measured at the specimen position at the lowest intensity level (×1) and converted to PFD. The ×1 level was set to approximately equal quanta across wavelengths (mean 1.04 × 10¹¹ photons cm⁻² s⁻¹, equivalent to 1.72 nmol m⁻² s⁻¹; range 0.93–1.13 × 10¹¹, CV 6.9%). Higher levels were obtained by removing neutral-density filters, giving nominal 3– and 9-fold intensities; PFD (×3) and PFD (×9) are calculated values. PFD is given in photons cm⁻² s⁻¹.

| wavelength(nm) | irradiance(nW/cm2) | PFD($\times 1$) | PFD($\times 3$) | PFD($\times 9$) |
| --- | --- | --- | --- | --- |
| 350 | 53.0 | 9.34E+10 | 2.801E+11 | 8.404E+11 |
| 400 | 51.0 | 1.03E+11 | 3.081E+11 | 9.243E+11 |
| 450 | 44.0 | 9.97E+10 | 2.99E+11 | 8.971E+11 |
| 500 | 43.7 | 1.10E+11 | 3.3E+11 | 9.9E+11 |
| 550 | 37.0 | 1.02E+11 | 3.073E+11 | 9.22E+11 |
| 600 | 36.0 | 1.09E+11 | 3.262E+11 | 9.786E+11 |
| 650 | 28.6 | 9.36E+10 | 2.808E+11 | 8.423E+11 |
| 700 | 31.0 | 1.09E+11 | 3.277E+11 | 9.832E+11 |
| 750 | 30.0 | 1.13E+11 | 3.398E+11 | 1.019E+12 |

**Table S4.**
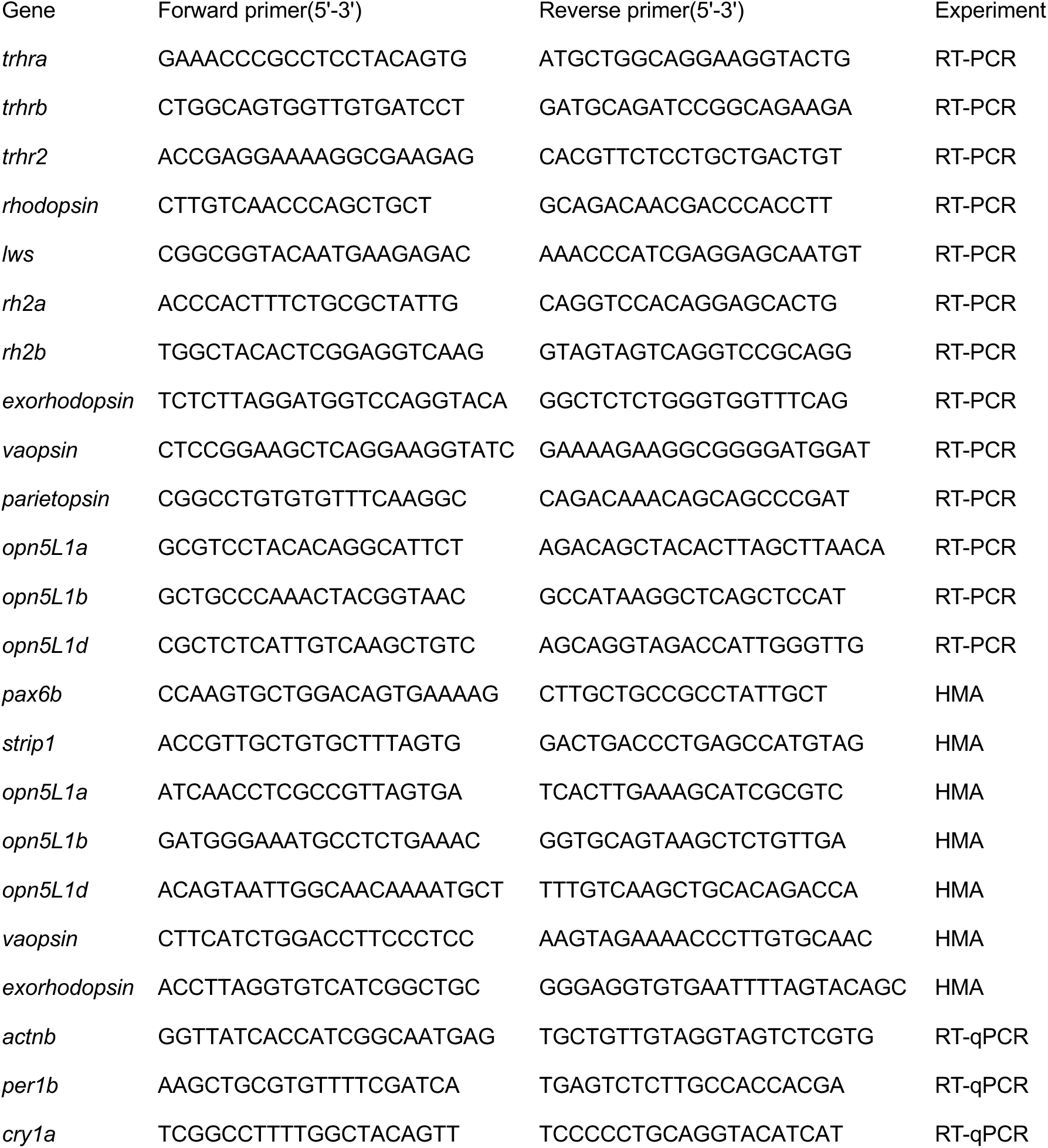
Forward and reverse primers for *trh* receptor genes, opsin genes and clock genes, used for RT-PCR, RT-qPCR and HMA.

## References

1. Ahmed, M., Kojima, Y., Masai, I. (2022) Strip1 regulates retinal ganglion cell survival by suppressing Jun-mediated apoptosis to promote retinal neural circuit formation. eLife 11, e74650. doi:10.7554/eLife.74650

2. Chen, T.W., Wardill, T.J., Sun, Y., Pulver, S.R., Renninger, S.L., Baohan, A., Schreiter, E.R., Kerr, R.A., Orger, M.B., Jayaraman, V., Looger, L.L., Svoboda, K., Kim, D.S. (2013) Ultrasensitive fluorescent proteins for imaging neuronal activity. Nature 499, 295–300. doi:10.1038/nature12354

3. Chinen, A., Hamaoka, T., Yamada, Y., Kawamura, S. (2003) Gene duplication and spectral diversification of cone visual pigments of zebrafish. Genetics 163, 663–675. doi:10.1093/genetics/163.2.663

4. Dimichele, L., Taylor, M.H. (1980) The environmental control of hatching in *Fundulus heteroclitus*. J. Exp. Zool. 214, 181–187. doi:10.1002/jez.1402140209

5. Dimichele, L., Taylor, M.H. (1981) The mechanism of hatching in *Fundulus heteroclitus*: development and physiology. J. Exp. Zool. 217, 73–79. doi:10.1002/jez.1402170108

6. Fobert, E.K., Burke da Silva, K., Swearer, S.E. (2019) Artificial light at night causes reproductive failure in clownfish. Biol. Lett. 15, 20190272. doi:10.1098/rsbl.2019.0272

7. Fobert, E. K., Schubert, K. P., Burke da Silva, K. (2021) The influence of spectral composition of artificial light at night on clownfish reproductive success. J. Exp. Mar. Biol. Ecol. 540, 151559. doi:10.1016/j.jembe.2021.151559

8. Forsell, J., Holmqvist, B., Helvik, J.V., Ekström, P. (1997) Role of the pineal organ in the photoregulated hatching of the Atlantic halibut. Int. J. Dev. Biol. 41, 591–595.

9. Fukuda, A., Sato, K., Fujimori, C., Yamashita, T., Takeuchi, A., Ohuchi, H., Umatani, C., Kanda, S. (2025) Direct photoreception by pituitary endocrine cells regulates hormone release and pigmentation. Science 387, 43–48. doi:10.1126/science.adj9687

10. Gajbhiye, D.S., Fernandes, G.L., Oz, I., Nahmias, Y., Golan, M. (2024) A transient neurohormonal circuit controls hatching in fish. Science 386, 1173–1178. doi:10.1126/science.ado8929

11. García-Fernández, J.M., Cernuda-Cernuda, R., Davies, W.I.L., Rodgers, J., Turton, M., Peirson, S.N., Follett, B.K., Halford, S., Hughes, S., Hankins, M.W., Foster, R.G. (2015) The hypothalamic photoreceptors regulating seasonal reproduction in birds: a prime role for VA opsin. Front. Neuroendocrinol. 37, 13–28. doi:10.1016/j.yfrne.2014.11.001

12. Gladstone, W. (2007) Temporal patterns of spawning and hatching in a spawning aggregation of the temperate reef fish Chromis hypsilepis (Pomacentridae). Mar. Biol. 151, 1143–1152.

13. Griem, J.N., Martin, K.L.M. (2000) Wave action: the environmental trigger for hatching in the California grunion *Leuresthes tenuis* (Teleostei: Atherinopsidae). Mar. Biol. 137, 177–181. doi:10.1007/s002270000329

14. Helvik, J.V., Walther, B.T. (1992) Photo-regulation of the hatching process of halibut (*Hippoglossus hippoglossus*) eggs. J. Exp. Zool. 263, 204–209. doi:10.1002/jez.1402630210

15. Helvik, J.V., Walther, B.T. (1993) Environmental parameters affecting induction of hatching in halibut (*Hippoglossus hippoglossus*) embryos. Mar. Biol. 116, 39–45.

16. Hinkle, P.M., Gehret, A.U., Jones, B.W. (2012) Desensitization, trafficking, and resensitization of the pituitary thyrotropin-releasing hormone receptor. Front. Neurosci. 6, 180. doi:10.3389/fnins.2012.00180

17. Kawaguchi, M., Yasumasu, S., Shimizu, A., Kudo, N., Sano, K., Iuchi, I., Nishida, M. (2013) Adaptive evolution of fish hatching enzyme: one amino acid substitution results in differential salt dependency of the enzyme. J. Exp. Biol. 216, 1609–1615. doi:10.1242/jeb.069716

18. Kingsford, M.J. (1985) The demersal eggs and planktonic larvae of *Chromis dispilus* (Teleostei: Pomacentridae) in north-eastern New Zealand coastal waters. N. Z. J. Mar. Freshw. Res. 19, 429–438. doi:10.1080/00288330.1985.9516107

19. Kleinjan, D.A., Bancewicz, R.M., Gautier, P., Dahm, R., Schonthaler, H.B., Damante, G., Seawright, A., Hever, A.M., Yeyati, P.L., van Heyningen, V., Coutinho, P. (2008) Subfunctionalization of duplicated zebrafish *pax6* genes by cis-regulatory divergence. PLoS Genet. 4, e29. doi:10.1371/journal.pgen.0040029

20. Kojima, D., Torii, M., Fukada, Y., Dowling, J.E. (2008) Differential expression of duplicated VAL-opsin genes in the developing zebrafish. J. Neurochem. 104, 1364– 1371. doi:10.1111/j.1471-4159.2007.05093.x

21. Lin, C.-H., Takahashi, S., Mulla, A.J., Nozawa, Y. (2021) Moonrise timing is key for synchronized spawning in coral *Dipsastraea speciosa*. Proc. Natl. Acad. Sci. USA 118, e2101985118. doi:10.1073/pnas.2101985118

22. Martin, K., Bailey, K., Moravek, C., Carlson, K. (2011) Taking the plunge: California grunion embryos emerge rapidly with environmentally cued hatching. Integr. Comp. Biol. 51, 26–37. doi:10.1093/icb/icr037

23. Matsumoto, Y., Fukamachi, S., Mitani, H., Kawamura, S. (2006) Functional characterization of visual opsin repertoire in Medaka (*Oryzias latipes*). Gene 371, 268–278. doi:10.1016/j.gene.2005.12.005

24. McAlary, F.A., McFarland, W.N. (1993) The effect of light and darkness on hatching in the pomacentrid *Abudefduf saxatilis*. Environ. Biol. Fishes 37, 237–244. doi:10.1007/BF00004631

25. Mitchell, L.J., Tettamanti, V., Rhodes, J.S., Marshall, N.J., Cheney, K.L., Cortesi, F. (2021) CRISPR/Cas9-mediated generation of biallelic F0 anemonefish (*Amphiprion ocellaris*) mutants. PLoS One 16, e0261331. doi:10.1371/journal.pone.0261331

26. Quiroga Artigas, G., Lapébie, P., Leclère, L., Takeda, N., Deguchi, R., Jékely, G., Momose, T., Houliston, E. (2018) A gonad-expressed opsin mediates light-induced spawning in the jellyfish *Clytia*. eLife 7, e29555. doi:10.7554/eLife.29555

27. Salis, P., Lee, S.-h., Roux, N., Lecchini, D., Laudet, V. (2021) The real Nemo movie: description of embryonic development in *Amphiprion ocellaris* from first division to hatching. Dev. Dyn. 250, 1651–1667. doi:10.1002/dvdy.354

28. Sato, K., Yamashita, T., Ohuchi, H., Takeuchi, A., Gotoh, H., Ono, K., Mizuno, M., Mizutani, Y., Tomonari, S., Sakai, K., Imamoto, Y., Wada, A., Shichida, Y. (2018) Opn5L1 is a retinal receptor that behaves as a reverse and self-regenerating photoreceptor. Nat. Commun. 9, 1255. doi:10.1038/s41467-018-03603-3

29. Schoots, A.F.M., Sackers, R.J., de Bont, R.G., Denucé, J.M. (1981) Ionophore A23187-induced hatching enzyme secretion in medaka embryos. Arch. Int. Physiol. Biochim. 89, B77–B78.

30. Smyder, E.A., Martin, K.L.M. (2002) Temperature effects on egg survival and hatching during the extended incubation period of California grunion, *Leuresthes tenuis*. Copeia 2002, 313–320. doi:10.1643/0045-8511(2002)002[0313:TEOESA]2.0.CO;2

31. Sweeney, A.M., Boch, C.A., Johnsen, S., Morse, D.E. (2011) Twilight spectral dynamics and the coral reef invertebrate spawning response. J. Exp. Biol. 214, 770–777. doi:10.1242/jeb.043406

32. Takeda, N., Kon, Y., Quiroga Artigas, G., Lapébie, P., Barreau, C., Koizumi, O., Kishimoto, T., Tachibana, K., Houliston, E., Deguchi, R. (2018) Identification of jellyfish neuropeptides that act directly as oocyte maturation-inducing hormones. Development 145, dev156786. doi:10.1242/dev.156786

33. Tarttelin, E.E., Fransen, M.P., Edwards, P.C., Hankins, M.W., Schertler, G.F.X., Vogel, R., Lucas, R.J., Bellingham, J. (2011) Adaptation of pineal expressed teleost exo-rod opsin to non-image forming photoreception through enhanced Meta II decay. Cell. Mol. Life Sci. 68, 3713–3723. doi:10.1007/s00018-011-0665-y

34. Wada, S., Shen, B., Kawano-Yamashita, E., Nagata, T., Hibi, M., Tamotsu, S., Koyanagi, M., Terakita, A. (2018) Color opponency with a single kind of bistable opsin in the zebrafish pineal organ. Proc. Natl. Acad. Sci. USA 115, 11310–11315. doi:10.1073/pnas.1802592115

35. Watanabe, M., Furuya, M., Miyoshi, Y., Inoue, Y., Iwahashi, I., Matsumoto, K. (1982) Design and performance of the Okazaki Large Spectrograph for photobiological research. Photochem. Photobiol. 36, 491–498. doi:10.1111/j.1751-1097.1982.tb04407.x

36. Yamagami, K. (1988) Mechanisms of hatching in fish. In Fish Physiology, Vol. 11A (eds Hoar, W.S., Randall, D.J.), pp. 447–499. Academic Press, San Diego.

37. Yamanaka, S., Kawaguchi, M., Yasumasu, S., Sato, K., Kinoshita, M. (2025) Effects of light and water agitation on hatching processes in false clownfish *Amphiprion ocellaris*. J. Exp. Zool. B Mol. Dev. Evol. 344, 29–40. doi:10.1002/jez.b.23276

38. Yamanaka, S., Okada, Y., Furuta, T., Kinoshita, M. (2021) Establishment of culture and microinjection methods for false clownfish embryos without parental care. Dev. Growth Differ. 63, 459–466. doi:10.1111/dgd.12759

39. Yasumasu, S., Katow, S., Umino, Y., Iuchi, I., Yamagami, K. (1989) A unique proteolytic action of HCE, a constituent protease of a fish hatching enzyme: tight binding to its natural substrate, egg envelope. Biochem. Biophys. Res. Commun. 162, 58–63. doi:10.1016/0006-291X(89)91961-X

40. Yasumasu, S., Kawaguchi, M., Ouchi, S., Sano, K., Murata, K., Sugiyama, H., Akema, T., Iuchi, I. (2010) Mechanism of egg envelope digestion by hatching enzymes, HCE and LCE in medaka, *Oryzias latipes*. J. Biochem. 148, 439–448. doi:10.1093/jb/mvq086

## References

41. Capella-Gutiérrez, S., Silla-Martínez, J. M., & Gabaldón, T. (2009). trimAl: A tool for automated alignment trimming in large-scale phylogenetic analyses. Bioinformatics, 25, 1972–1973.

42. Chen, S., Zhou, Y., Chen, Y., Gu, J. (2018) fastp: an ultra-fast all-in-one FASTQ preprocessor. Bioinformatics 34, i884–i890. doi:10.1093/bioinformatics/bty560

43. Dobin, A., Davis, C.A., Schlesinger, F., Drenkow, J., Zaleski, C., Jha, S., Batut, P., Chaisson, M., Gingeras, T.R. (2013) STAR: ultrafast universal RNA-seq aligner. Bioinformatics 29, 15–21. doi:10.1093/bioinformatics/bts635

44. Durinck, S., Spellman, P.T., Birney, E., Huber, W. (2009) Mapping identifiers for the integration of genomic datasets with the R/Bioconductor package biomaRt. Nat. Protoc. 4, 1184–1191. doi:10.1038/nprot.2009.97

45. Katoh, K., & Standley, D. M. (2013). MAFFT multiple sequence alignment software version 7: Improvements in performance and usability. Molecular Biology and Evolution, 30, 772–780.

46. Li, B., Dewey, C.N. (2011) RSEM: accurate transcript quantification from RNA-Seq data with or without a reference genome. BMC Bioinformatics 12, 323. doi:10.1186/1471-2105-12-323

47. Robinson, M.D., McCarthy, D.J., Smyth, G.K. (2010) edgeR: a Bioconductor package for differential expression analysis of digital gene expression data. Bioinformatics 26, 139–140. doi:10.1093/bioinformatics/btp616

48. Stamatakis, A. (2014). RAxML version 8: A tool for phylogenetic analysis and post-analysis of large phylogenies. Bioinformatics, 30, 1312–1313.

49. Whelan, S., & Goldman, N. (2001). A general empirical model of protein evolution derived from multiple protein families using a maximum-likelihood approach. Molecular Biology and Evolution, 18, 691–699.

